# BR-bodies link RNA homeostasis to stress tolerance and intracellular fitness in *Brucella*

**DOI:** 10.64898/2026.07.11.737834

**Authors:** Kaveendya S. Mallikaarachchi, Rosemary Northcote, Thomas Kim, Melene A. Alakavuklar, Vincent Lawal, Vidhyadhar Nandana, Aretha Fiebig, Jared M. Schrader, Sean Crosson

## Abstract

Bacterial gene expression depends on the coordinated regulation of RNA synthesis, processing, and decay. The conserved RNA degradosome scaffold, Ribonuclease E (RNase E), assembles into phase-separated bacterial ribonucleoprotein bodies (BR-bodies) through its C-terminal intrinsically disordered region (IDR). This IDR also scaffolds recruitment of degradosome client proteins that carry out post-transcriptional gene regulation. Whether BR-bodies regulate virulence programs to support infection, however, is largely undefined. We show that RNase E of the intracellular pathogen *Brucella ovis* forms RNA-dependent condensates in vivo and phase-separates with RNA in vitro, with the IDR necessary and sufficient for BR-body assembly. Deleting the IDR (*rne*(ΔIDR)) did not impair growth but sensitized *Brucella* to host-relevant oxidative and cell-envelope stressors. Transcriptome-wide profiling that simultaneously resolved mRNA decay and processing revealed that BR-bodies primarily accelerate mRNA turnover while stabilizing a distinct subset of transcripts. Notably, BR-bodies control the processing and levels of *virB* type IV secretion system (T4SS) mRNA and the turnover of its key activators, linking condensate-based RNA regulation to a core virulence pathway. Consistent with *virB* dysregulation, the *rne*(ΔIDR) mutant was severely attenuated in mammalian macrophages. To test whether phase separation is sufficient for BR-body function, we replaced the *B. ovis* IDR with the highly divergent *Caulobacter crescentus* IDR. This chimera assembled BR-bodies but rescued fitness incompletely, fully restoring oxidative-stress resistance but not cell-envelope stress resistance or intracellular fitness. BR-bodies therefore link RNA metabolism to *Brucella* stress resistance and infection, and their full function requires both phase separation and additional native IDR-specific activities such as degradosome interactions.

**Importance:** Biomolecular condensates are non-membrane bound organelles that organize biological processes across all domains of life, but their role in bacterial infection is not well characterized. We show that *Brucella*, an intracellular bacterial pathogen and causative agent of the disease brucellosis, relies on biomolecular condensates called BR-bodies to control RNA stability and regulate gene expression. Without BR-bodies, *Brucella* is sensitized to host-relevant chemical stresses, cannot properly regulate essential infection machinery, and has severely diminished fitness within mammalian host cells. These results indicate that control of RNA levels by biomolecular condensates is required for *Brucella* survival within animal hosts, providing evidence that phase separation is a fundamental mechanism underlying *Brucella* adaptation to the hostile environment encountered during infection.

## Introduction

To adapt to changing environments, cells must precisely balance the synthesis, processing, and decay of RNA that together determine gene expression. Biomolecular condensates have emerged as membraneless compartments that organize RNA metabolism through liquid-liquid phase separation (LLPS) of proteins and nucleic acids (1, 2). By concentrating specific factors and reactions, these dynamic structures help cells remodel their physiology in response to environmental change (3, 4). In eukaryotes, mRNA decay factors partition into stress granules and processing bodies (P-bodies), which are ribonucleoprotein (RNP) condensates whose formation reflects the translational state of the cell and promotes selective mRNA turnover and storage (5–7). Recent work has established that bacterial RNA metabolism is similarly structured by biomolecular condensation (8).

RNase E, the conserved scaffold of the bacterial RNA degradosome (9–11), assembles into phase-separated bacterial ribonucleoprotein bodies (BR-bodies) that concentrate RNAs and decay factors to promote RNA turnover and control gene regulation (12–15). BR-body assembly is driven by the large, intrinsically disordered C-terminal region (IDR) of RNase E, which is both necessary and sufficient for phase separation and for recruitment of degradosome client proteins (12, 14). Although RNase E IDRs are poorly conserved at the sequence level, their broad phylogenetic distribution points to a conserved biophysical role in organizing RNA decay (8, 16, 17). Notably, distinct RNases play analogous condensate-organizing roles in other bacterial lineages: RNase Y forms dynamic foci in many Firmicutes (18–21), RNase J organizes peripheral degradosome structures in *Helicobacter pylori* (22), and PNPase organizes RNA degradosomes in human mitochondria (23). Their independent evolution across diverse phyla suggests that biomolecular condensation is an ancient widespread mechanism for organizing bacterial RNA homeostasis (3).

Disrupting RNase E condensation has important functional consequences. In α-proteobacteria, deletion of the IDR (*rne*(ΔIDR)) alters global mRNA decay (13) and impairs fitness under stress in *Sinorhizobium meliloti* (24) and *Caulobacter crescentus* (25). These results indicate that BR-bodies are adaptive regulators linking transcript fate with physiological state. More recently, BR-bodies have been shown to do more than support basal mRNA decay. Specifically, they can reorganize into a solid-like state under stress, shifting toward sequestration and protection of mRNAs from decay (26). More broadly, transposon screens across γ- and β-proteobacterial pathogens (including *Vibrio*, *Pseudomonas*, *Salmonella*, *Neisseria*, and *Snodgrassella*) have repeatedly recovered C-terminal insertions in the RNase E gene that disrupt the IDR and attenuate host colonization, hinting that BR-body-mediated RNA regulation may generally support infection (27–33).

*Brucella* species are intracellular α-proteobacterial pathogens that cause reproductive failure in livestock and zoonotic disease in humans worldwide (34, 35). Because survival within host phagocytes requires rapid reprogramming of gene expression, regulated RNA metabolism is likely central to *Brucella* infection. Supporting this, a *Brucella abortus* mutant lacking the RNase E C-terminus shows reduced spleen colonization in a mouse model of infection and dysregulated levels of nearly 300 transcripts (36). These results implicate the *Brucella* RNase E IDR in coordinating expression of genes required for infection and colonization. However, it remains unknown whether the IDR drives BR-body formation in *Brucella* and how BR-bodies contribute to stress resilience and fitness in the intracellular niche.

To address these questions, we generated an RNase E variant lacking the IDR (*rne*(ΔIDR)) in the ovine intracellular pathogen *Brucella ovis*. We establish that the IDR is required for BR-body assembly in the *B. ovis* cytosol and for RNase E phase separation *in vitro*. We used a transcriptome-wide assay of RNA decay and processing to link BR-bodies to the regulation of key transcripts required for host infection, including the *virB* type IV secretion system (T4SS) and its regulators. To test whether condensation alone accounts for BR-body function, we replaced the native IDR with the highly divergent *Caulobacter crescentus* IDR, generating a chimeric RNase E (*rne*(IDR_Cc_)) that phase-separates but is predicted to have reduced interactions with *Brucella* degradosome clients. Comparing this chimera with wild-type and IDR-deletion strains across stress and macrophage-infection assays permitted us to separate the physical process of phase separation from the lineage-specific interactions encoded by the IDR.

Together, these experiments position *Brucella* BR-bodies at the interface of RNA metabolism and intracellular pathogenesis and indicate that phase separation is necessary, but not sufficient, for full BR-body function. By linking BR-body assembly to the coordinated control of the T4SS and stress resistance, this study shows how biomolecular condensates support adaptive physiology and effector-mediated infection programs during *Brucella* colonization of mammalian phagocytes.

## Materials and Methods

### Bacterial cell growth

*Brucella ovis* ATCC 25840 and derivative strains were cultured on tryptic soy agar (TSA; BD Difco) plates supplemented with 5% sheep blood (Quad Five), or in Brucella broth. All strains were grown at 37°C with 5% CO_2_ supplementation, except for strains harboring a functional carbonic anhydrase (*bcaA1*) (37) which can be grown at 37°C without supplemented CO_2_.

*Escherichia coli* strains used for generating mutants were all grown in lysogeny broth (LB), or LB solidified with 1.5% wt/vol agar. Both Top 10 and WM3064 cultures were grown at 37°C, with 500rpm shaking for liquid cultures. WM3064 (gift of William Metcalf) was grown with 30 µM diaminopimelic acid (DAP) supplementation. Media was supplemented with 50 µg ml^−1^ kanamycin when necessary for plasmid maintenance. Primer, plasmid, and strain information can be found in Table S5. Bacterial growth curves were carried out in a Tecan Infinite 200 Pro Plate reader, monitoring at 600 nm.

### Plasmid and Strain Construction

#### Chromosomal deletion and complementation strain construction

All *B. ovis* strains harboring in-frame, unmarked deletions were generated using double-crossover recombination. Briefly, approximately 500 base pairs (bp) up and downstream of the target region were amplified via PCR from template *B. ovis* genomic DNA using KOD Xtreme polymerase (Novagen) with primers listed in Table S1. Regions up and downstream were stitched together with an overlapping PCR using external primers and the resulting DNA fragment was inserted into pNPTS138, a *sacB*-containing suicide plasmid, using restriction enzyme digestion and ligation. Generated plasmids were chemically transformed into *E. coli* Top 10 and confirmed via sequencing. Plasmids were then transformed into WM3064, a donor strain of *E. coli* that is a DAP auxotroph and conjugation competent. The desired plasmid was transferred from WM3064 to *B. ovis* via conjugation. *Brucella* cells were plated on TSA blood plates with 50µg/ml kanamycin to select for primary recombinants, then grown overnight in non-selective brucella broth. Overnight cultures were plated on 5% sucrose plates to identify clones that underwent a second recombination event to remove the plasmid. Kanamycin sensitive colonies were screened via colony PCR to distinguish deletion mutants from wild type. The *C. crescentus* RNase E chimera was made in a similar manner, amplifying ∼250 bp up and downstream of the IDR domain in *B. ovis*, as well as the IDR domain of RNase E in *C. crescentus* NA1000. The three fragments were used in two overlapping PCRs, the first stitching together the downstream region of *Brucella* RNase E with the IDR domain of *Caulobacter*, then adding the upstream region from *B. ovis* to the front to create the insert. For all complementation strains, the insert was generated by amplifying target regions plus ∼250 bp up and downstream to restore the genes to the native locus. All subsequent steps for the double-recombination method were followed as described above. All strains with a functional carbonic anhydrase (*bcaA1*) (which permits growth without supplemented CO_2_) were generated in the same manner as described above, except the plasmids were mated into a *B. ovis* strain harboring the *bcaA1* allele described by Varesio *et. al* (37).

#### Fluorescent RNase E strain construction

*B. ovis bcaA1 rne-*GFP and *rne*(ΔIDR)-GFP strains were also constructed using double-crossover recombination, adding a C-terminal GFP tag to RNase E with (*rne-*GFP) or without (*rne*(ΔIDR)-GFP) the IDR domain at the native locus. Approximately 500bp up- and downstream of the 3’ end of *rne* or *rne*(ΔIDR) was amplified via PCR as described above with primers listed in Table S4. Monomeric superfolder green fluorescent protein (msfGFP, henceforth GFP) was amplified from a plasmid, and the three fragments were used in two overlapping PCRs. The first PCR stitched together the upstream region with GFP, and the second stitched the upstream and GFP fragment with the downstream region. The generated fragment was inserted into pNPTS138 and all subsequent steps for the double-recombination method were followed as described above. For the *rne*(IDR_Cc_)-GFP strain, GFP and the downstream region of *rne* was amplified from the pNTPS138-*rne*-GFP plasmid generated above via PCR, as well as the upstream region of the chimeric *rne* gene from *rne*(IDR_Cc_) genomic DNA. The two fragments were used in an overlapping PCR, inserted into pNTPS138, and again all subsequent steps for the double-recombination method were followed as described. Strains with the dsRed background were generated by integration of a pUC18-mTn7K-dsRed.M1 plasmid containing the fluorescent reporter into the *glmS* locus (38). The plasmid was co-conjugated with pTNS3 (Addgene), a suicide helper plasmid expressing the Tn7 transposase. *B. ovis* colonies carrying the integrated mTn7K-dsRed.M1 at the *glmS* locus were selected for on TSA blood plates with 50µg/ml kanamycin and checked for fluorescence intensity.

#### Plasmid construction for recombinant protein expression

The pET His₆-MBP-TEV LIC cloning vector (1M) was used as the backbone to generate the pKM2 (encoding full-length RNase E), pKM42 (encoding RNase EΔIDR, corresponding to the nuclease domain only), and pKM43 (encoding the RNase E intrinsically disordered region [IDR] only) plasmids for recombinant protein expression in *E. coli*. All constructs encode N-terminally 6×His-MBP-tagged proteins under the control of an IPTG-inducible T7 promoter. The pET His₆-MBP-TEV LIC cloning vector (1M) was a gift from Scott Gradia (Addgene plasmid #29656; RRID:Addgene_29656).

To generate pKM2, the *rne* coding sequence was PCR-amplified from *B. ovis* ATCC 25840 genomic DNA. The pET His₆-MBP-TEV LIC vector was linearized with SspI, and the amplified *rne* fragment was inserted by ligation-independent cloning. The resulting construct was transformed into *E. coli* DH5α cells. Kanamycin-resistant colonies were grown overnight in LB medium, plasmids were isolated by miniprep, screened by restriction enzyme digestion, and verified by whole-plasmid sequencing (Genewiz).

The pKM42 and pKM43 plasmids were generated from pKM2 by inverse PCR. Following PCR amplification, the parental template was digested with DpnI, and the PCR product was purified after agarose gel electrophoresis, ligated using T4 DNA ligase (Thermo Scientific), and transformed into *E. coli* DH5α cells. Kanamycin-resistant colonies were grown overnight in LB medium, plasmids were isolated by miniprep, screened by restriction enzyme digestion, and verified by whole-plasmid sequencing (Genewiz).

#### Cell Imaging experiments

For imaging, cells from exponential-phase cultures (OD_600_ 0.3-0.5) were immobilized on 1.5% agarose pads prepared with Brucella broth and mounted on microscope slides. Images were acquired using a Nikon Eclipse T*i*2 microscope equipped with a Prime BSI SCMOS camera and a 100× oil immersion CFL Plan Fluor DLL objective (Nikon), controlled by NIS-Elements software. The ET-GFP filter set was used for GFP imaging (Nikon 96362), and the ET-mCH/TR filter set was used for dsRed imaging (Nikon 96365).

#### Image analysis

Foci per cell were quantified manually by counting the number of cells and foci from three independent biological replicates for each condition. Fluorescence intensity and foci area in time-lapse images were measured using Fiji (ImageJ). Individual foci were manually outlined, and the area and total fluorescence within each focus were measured.

To determine the fraction of fluorescence localized within foci, cells were outlined using the Cell tool in Fiji (ImageJ), and the cell area and total cellular integrated fluorescence intensity were measured. Foci within each cell were then manually outlined, and the area and integrated fluorescence intensity of each focus were measured. The fraction of fluorescence localized within foci was calculated by dividing the summed integrated fluorescence intensity of all foci in a cell by the total integrated cellular fluorescence intensity.

For line trace analysis, a straight line was drawn along the length of each cell, and GFP and dsRed fluorescence intensity profiles were extracted using Fiji (ImageJ). For correlation coefficient analysis, line profiles were generated as described above, and fluorescence intensities from the GFP and dsRed channels were measured at corresponding positions along each line. Pearson’s correlation coefficients were calculated from the paired fluorescence intensity values for each cell.

#### Stress resistance/tolerance experiments

*B. ovis* strains (wild type 25840, *rne*(ΔIDR), *rne*^+^, and *rne*(IDR_Cc_)) were resuspended in phosphate buffered saline (PBS) from 48–72-hour old TSA blood plates. Strains were normalized to an OD_600_ of 0.3 and five microliters of each dilution in a log_10_ dilution series was plated on TSAB or TSAB supplemented with 1.5ug/ml carbenicillin or 8ug/ml polymyxin B. For the oxidative stress assay, strains were resuspended as described above but normalized to an OD_600_ of 0.15. Cultures were then resuspended in PBS mixed with hydrogen peroxide to a final concentration of 0.3% H_2_O_2_, incubated at 37°C for 30 minutes, then serially diluted and plated on TSAB as described above. Plates were incubated at 37°C + 5% CO_2_, and images of titer plates were taken after 72-96 hours.

#### Rif-seq

To measure transcript half-lives and map RNA ends genome-wide, we performed rifampicin sequencing (Rif-seq) using end-enriched RNA-sequencing (REND-seq (39)) libraries. *B. ovis* strains (wild type 25840 and *rne*(ΔIDR)) were grown in Brucella broth overnight. The next day, exponential-phase cultures (OD_600_ 0.3–0.5) were treated with 10 mg/mL rifampicin to achieve a final concentration of 200 µg/mL. Before adding rifampicin, 1 mL of culture (0 minutes) was removed and mixed with 2 mL of RNAprotect Bacterial Reagent (QIAGEN), vortexed, and incubated at room temperature for 5 minutes. This step was repeated at 1, 3, and 9 minutes after rifampicin was added. Then, the cells were pelleted at 20,000 × *g* for 1 minute and resuspended in 1 mL of 65°C TRIzol Reagent (Ambion) and incubated at 65°C for 10 minutes in a heat block. 200 µL of chloroform was added to the samples, and the tubes were incubated at room temperature for 5 minutes before spinning at 20,000 × *g* for 10 minutes. RNA samples were chloroform extracted once and precipitated using isopropanol (1× volume isopropanol, 0.1× volume 5 M sodium acetate, pH 5.2, 2 µL of Glycoblue (ThermoFisher)) overnight at −80°C. The RNA samples were spun at 20,000 × *g* at 4°C for 1 hour; the pellets were washed with 80% ethanol for 10 minutes, air dried, and resuspended in 10 mM Tris-HCl (pH 7.0). The RNA-Seq libraries were made using 100 ng of total RNA samples, and end-enriched library construction was performed according to published protocols (39, 40). Reads were trimmed and aligned to the *B. ovis* ATCC 25840 genome (NC_009505; NC_009504). To measure RNA-decay rates, the fraction of mRNA remaining was calculated as the RPKM of each time point divided by the RPKM measured in the untreated 0 min sample. For bulk mRNA half-life calculations of RNA-seq data, the fraction of all mRNA reads was compared with the total fraction of reads, which includes a majority of stable tRNA reads. Half-lives of individual mRNAs were calculated using the Rifcorrect software package (41). All samples were uploaded to NCBI GEO under accession number GSE337175.

For steady state mRNA level analysis, the “0 minute” time point before rifampicin addition was compared for wild type and *rne*(ΔIDR) samples. RPKM levels from these samples were compared to determine which mRNAs had relatively higher and lower levels due to *rne* IDR domain deletion at steady state. RNAs were filtered for low counts, before significance class (more abundant, less abundant, or not significant) was calculated from a Benjamini-Hochberg-adjusted p < 0.05 cutoff for significance.

#### Gene function enrichment analysis

Gene Ontology overrepresentation analysis of the RNA-seq results was performed using the RefSeq annotation of *Brucella ovis* ATCC 25840 (NC_009505 and NC_009504; chromosomes I and II, respectively) downloaded as GenBank flat files. Coding sequences (CDS features) were parsed. For each CDS, we extracted the original locus identifier (/old_locus_tag) together with the corresponding current gene name and product description. Gene Ontology (GO) annotations were obtained directly from the RefSeq-supplied qualifiers /GO_process, /GO_function, and /GO_component. From these qualifiers we parsed the GO identifier (e.g. GO:0006412) and its textual description, and recorded the associated ontology namespace (biological process, molecular function, or cellular component).

Differentially expressed genes between the wild-type strain and the *rne*(ΔIDR) mutant (as defined in the RNA-seq analysis described above) were separated into two gene sets: transcripts with significantly higher steady-state levels in the *rne*(ΔIDR) mutant and transcripts with significantly lower steady-state levels. Only genes with at least one GO annotation were retained for enrichment testing. The background “universe” for the analysis was defined as all CDSs in the *B. ovis* genome with at least one GO term (n=1938). For each gene set (higher and lower transcript levels analyzed separately), GO term overrepresentation was tested using a custom Python implementation of the one-sided hypergeometric test (equivalent to Fisher’s exact test for a 2×2 contingency table), following the general framework used in GO enrichment tools (42, 43). Resulting p-values were adjusted for multiple testing using the Benjamini-Hochberg procedure to control the false discovery rate.

#### Gene set enrichment analysis

Gene set enrichment analysis (GSEA) was performed using the Functional Analysis module of OmicsBox. Functional annotations were generated using the Blast2GO workflow implemented in OmicsBox. Sequence similarity searches were performed using NCBI BLAST, and protein domain information was obtained using InterProScan. Gene Ontology (GO) mapping associated homologous sequences and conserved domains with GO annotations derived from the Gene Ontology Consortium.

Differentially expressed genes identified by RNA-seq were compiled into a ranked list based on log_2_ fold change between the wild-type strain and the *rne*(ΔIDR) mutant, with positive values indicating upregulation and negative values indicating downregulation. This ranked list was provided as the rank file input for GSEA. The background gene universe was defined as all annotated coding sequences in the *B. ovis* genome (n = 2755).

Statistical significance of enrichment was evaluated using gene set permutations, with the number of permutations specified as 500. Enrichment scores were calculated using the standard running-sum statistic, in which the score increases when a gene in the ranked list belongs to a given GO term and decreases otherwise. The maximum deviation from zero was reported as the enrichment score. GO terms from the biological process, molecular function, and cellular component ontologies were tested. Detailed results were generated for the top-ranked GO terms, and extended results were obtained for all significantly enriched GO categories. Resulting p values were adjusted for multiple testing using the Benjamini–Hochberg procedure to control the false discovery rate.

#### Western blot

1 mL of exponential-phase cells (OD_600_ 0.3-0.5) from *B. ovis bcaA1, bcaA1 rne-*GFP, *bcaA1 rne*(ΔIDR)-GFP, and *bcaA1 rne*(IDR_Cc_)-GFP cultures were pelleted and resuspended in 250 μL of 1x Laemmli buffer for each 1.0 OD_600_ unit and boiled at 95°C for 5 minutes. Next, 15 μL from each sample was loaded in a 4–20% Bio-Rad stain-free TGX precast gel alongside 1 μL of Positope Control Protein (Thermofisher R90050) and electroporated at 150V for 60 minutes. Then, semi-dry transfer was done using the BioRad Trans-Blot Turbo transfer system and Trans-Blot Turbo midi 0.2 µm nitrocellulose transfer pack according to the manufacturer’s instructions. Next, the membrane was blocked for 1 hour using 25 mL of 3% BSA in TBST (20 mM Tris, pH 7.5, 150 mM NaCl, 0.1% Tween-20) at room temperature with gentle shaking. For primary antibody binding, the membrane was then submerged in 1:1000 dilution of the anti-GFP antibody (Millipore Sigma 11814460001) in the same blocking buffer and shaken gently overnight at 4°C. After washing the membrane 5 times with TBST for 10 minutes each time, the membrane was probed with 1:20000 goat anti-mouse IgG secondary antibody with HRP conjugate (Millipore Sigma 12-349) in blocking buffer. Next, secondary antibody incubation was done for 1 hour with gentle shaking at room temperature. The membrane was then washed 5 times, 10 minutes each time, with 1x TBST with gentle shaking. Following the wash, the blot was treated with SuperSignal™ West Pico PLUS chemiluminescent reagent (ThermoFisher 34577) for 5 minutes and imaged using the Bio-Rad ChemiDoc MP imaging system.

#### Macrophage cell culture and infection assay

THP-1 Macrophage like cells were grown in complete Roswell Park Memorial Institute 1640 medium (RPMI) supplemented with 2 mM glutamine and 10% Heat Inactivated Fetal Bovine Serum (HI FBS) to a maximum titer of 1 x 10^6^ cells/ml. 48 hours prior to infection, THP-1 macrophage like cells were differentiated using 50 ng/ml phorbol-myristate acetate (PMA) and seeded at a titer of 1 x 10^5^ cells per well into 96-well plates. *Brucella ovis* wild type 25840, *rne*(ΔIDR), *rne*^+^, Δ*virB2, rne*(ΔIDR) *ΔvirB2,* and *rne*(IDR_Cc_) strains were scraped from 48–72-hour old TSA blood plates and resuspended in RPMI supplemented with 2mM glutamine and 10% HI FBS. All bacterial cultures were normalized and added to tissue culture plates at a multiplicity of infection (MOI) of 100. Plates were centrifuged at 150 x g for 5 minutes and incubated for 1 hour at 37°C + 5% CO_2_. Media was then removed from the wells and replaced with RPMI + 25 μg/ml gentamycin. At 2, 24, and 48 hours post infection, THP-1 cells were washed with PBS and subsequently lysed with water. Released *Brucella* were serially diluted in a log_10_ dilution series and plated on TSA blood plates to enumerate colony forming units (CFUs).

#### RNase E full length and deletion constructs protein purifications

*E. coli* BL21(DE3) cells transformed with pKM2, pKM42 and pKM42 were cultured in 2 L of 2×LB medium to an OD_600_ of 0.6-0.7. Protein expression was induced with 0.5 mM IPTG at 30°C for 3.5 hours. Cells were harvested by centrifugation at 6,000 rpm for 10 minutes and resuspended in 30 mL lysis buffer (20 mM Tris-HCl, pH 7.4, 500 mM NaCl, 10% glycerol, 20 mM imidazole) supplemented with 40 µg/mL DNase I, 1 mM PMSF, and protease inhibitor cocktail (3 tablets). Cells were lysed on ice by sonication (50% amplitude, 7 seconds pulse on/10 seconds pulse off cycles, total 4 minutes). Lysates were clarified by centrifugation at 14,000 rpm for 45 minutes, and the supernatant was applied to a 10 mL Ni-NTA affinity column pre-equilibrated with lysis buffer. The column was sequentially washed with 100 mL lysis buffer, 100 mL high-salt buffer (20 mM Tris-HCl, pH 7.4, 1 M NaCl, 10% glycerol, 10 mM imidazole), and 100 mL wash buffer (20 mM Tris-HCl, pH 7.4, 150 mM NaCl, 10% glycerol, 10 mM imidazole). Protein was eluted using elution buffer (20 mM Tris-HCl, pH 7.4, 150 mM NaCl, 5% glycerol, 250 mM imidazole) in 5 mL fractions. Eluted fractions containing protein were pooled and concentrated to ∼10 mg/mL, followed by size-exclusion chromatography on a Superdex 200 Increase 10/300 GL column equilibrated in SEC buffer (20 mM Tris-HCl, pH 7.4, 250 mM NaCl, 2% glycerol, 1 mM DTT). Purified protein was concentrated to ∼10 mg/mL, flash-frozen, and stored at -80°C.

For MBP tag removal, 6×His-MBP-RNase E was incubated with TEV protease at a 20:1 (substrate:protease) molar ratio overnight at 4°C in SEC buffer. The cleaved sample was subjected to a second Superdex 200 Increase 10/300 GL run to separate RNase E from the cleaved tag. Purified RNase E was concentrated to ∼10 mg/mL and stored at -80°C.

#### *In vitro* phase separation assays

Phase separation assays were carried out using purified preparations of full length RNase E and MBP fusions of RNase E delta NTD (i.e. the nuclease domain) and delta CTD (i.e. the IDR).

Phase separation of full-length RNase E: 6 µM of *Brucella ovis* RNase E was incubated with 40 ng/µL *E. coli* total RNA in 20 mM Tris pH 7.4, 100 mM NaCl, 1 mM DTT buffer in a total reaction volume of 10 µl for 30 minutes at room temperature. Entire 10 µl was spotted on a slide and covered with coverslip before imaging with Nikon Eclipse NI-E with Prime BSI SCMOS camera and a 100× oil immersion CFL Plan Fluor DLL objective (Nikon) under phase contrast, with 30 millisecond exposure.

Phase separation of MBP-RNase EΔIDR or MBP-RNase E IDR only: 6 µM of *Brucella ovis* MBP-RNase EΔIDR or IDR only was incubated with or without 40 ng/µL *E. coli* total RNA, and 0.25 µM TEV protease in 20 mM Tris pH 7.4, 100 mM NaCl, 1 mM DTT buffer in a total reaction volume of 10 µl for 1 hour at room temperature. The 10 µl sample was spotted on a slide and covered with coverslip before imaging with Nikon Eclipse NI-E with Prime BSI SCMOS camera and a 100× oil immersion CFL Plan Fluor DLL objective (Nikon) under phase contrast, with 30 millisecond exposure.

## Results

### *Brucella* RNase E forms RNA-dependent BR-body condensates *in vivo* and *in vitro*

*B. ovis* RNase E contains a large, low-complexity intrinsically disordered region (IDR) appended to the C-terminus of the highly conserved catalytic domain (Fig. 1A-C); this architecture is shared among α-proteobacterial RNase E homologs where the IDR scaffolds bacterial ribonucleoprotein bodies (BR-bodies) (12, 24). This *B. ovis* RNase E IDR contains patches of positively and negatively charged amino acids (Fig. 1D) which were previously shown to drive RNase E condensation (12). Although α-proteobacterial IDRs share little primary structure identity and vary considerably in length, this charge patterning is broadly conserved, suggesting a shared biophysical mechanism for BR-body assembly across species (12).

**Figure 1.**
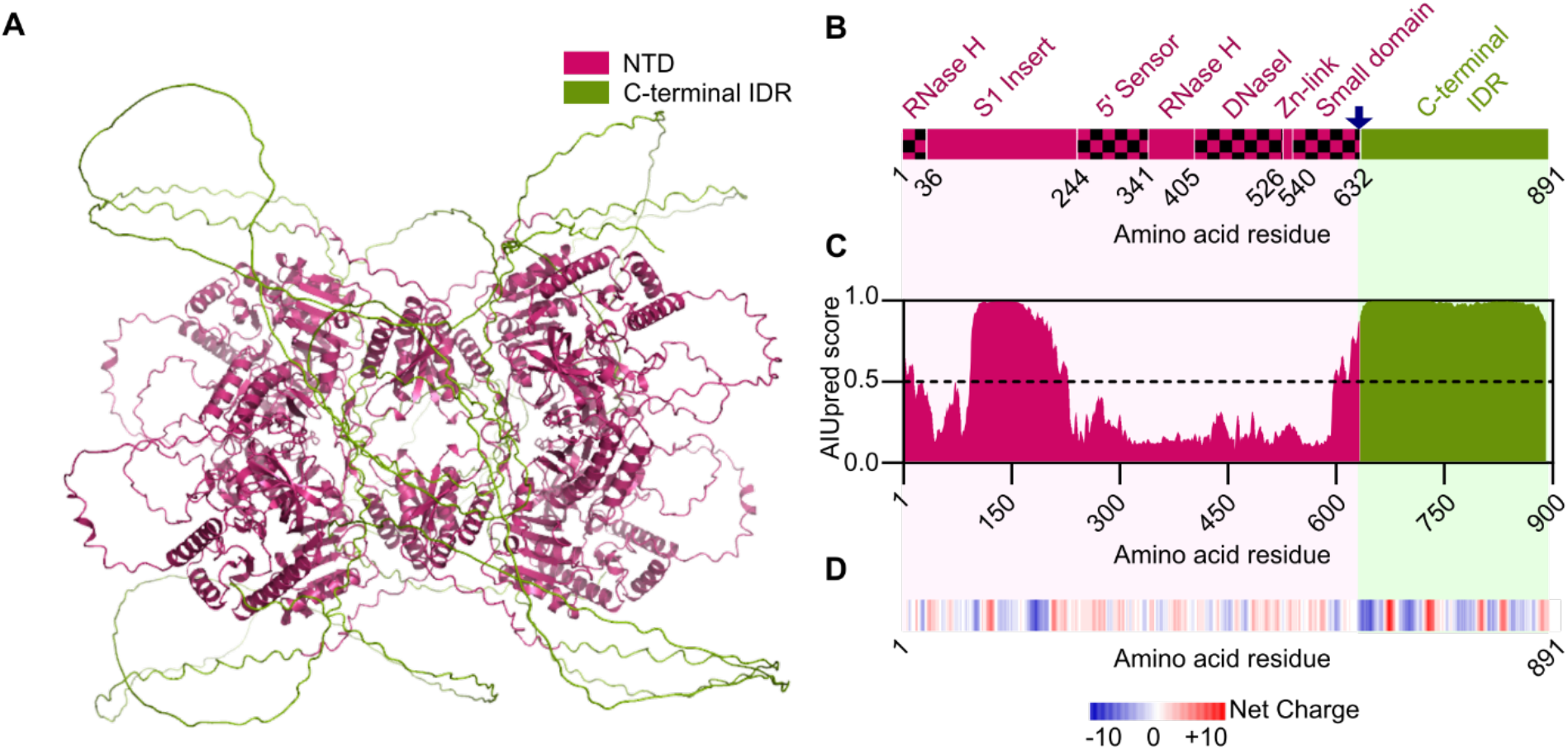
*Brucella ovis* RNase E contains sequence features necessary for phase separation. **A.)** Predicted AlphaFold3 structure of the *B. ovis* RNase E tetramer (68). The folded N-terminal catalytic nuclease domain is shown in pink while the C-terminal intrinsically disordered region (IDR) is shown in green. **B.)** Schematic of the *B. ovis* RNase E domain organization, with conserved motifs and domain architecture annotated. Domain boundaries were assigned based on Hardwick *et al*. (65). **C)** AIUPred disorder profile (69) for *B. ovis* RNase E. Values above 0.5 (dotted line) indicate residues predicted to be intrinsically disordered. **D)** Charge distribution across *B. ovis* RNase E calculated as net charge within a 10–amino acid sliding window. Positively charged residues are shown in red, and negatively charged residues are shown in blue.

To test whether RNase E forms BR-body condensates inside the *B. ovis* cell, we examined the localization of a functional RNase E- monomeric superfolder GFP fusion expressed from the native chromosomal locus (*rne*-GFP). In exponentially growing cells, RNase E-GFP formed discrete cytoplasmic foci rather than a diffuse signal (Fig. 2A; Fig. S1). Time-lapse live cell imaging revealed that these foci were dynamic, undergoing fusion events that produced larger condensates with increased fluorescence intensity, consistent with liquid-like behavior reported for other α-proteobacterial BR-bodies (3, 24) (Fig. 2B). To assess phase separation *in vitro,* we examined purified full-length *B. ovis* RNase E under physiological protein and salt conditions. RNase E remained soluble in the absence of RNA but condensed into spherical, micron-scale droplets upon RNA addition, consistent with RNA-driven phase separation (Fig. 2C). This RNA dependence mirrors observations in other α-proteobacteria (12, 24) supporting the conclusion that *B. ovis* RNase E forms RNA-dependent biomolecular condensates *in vitro* that likely correspond to the BR-body foci observed *in vivo*.

**Figure 2.**
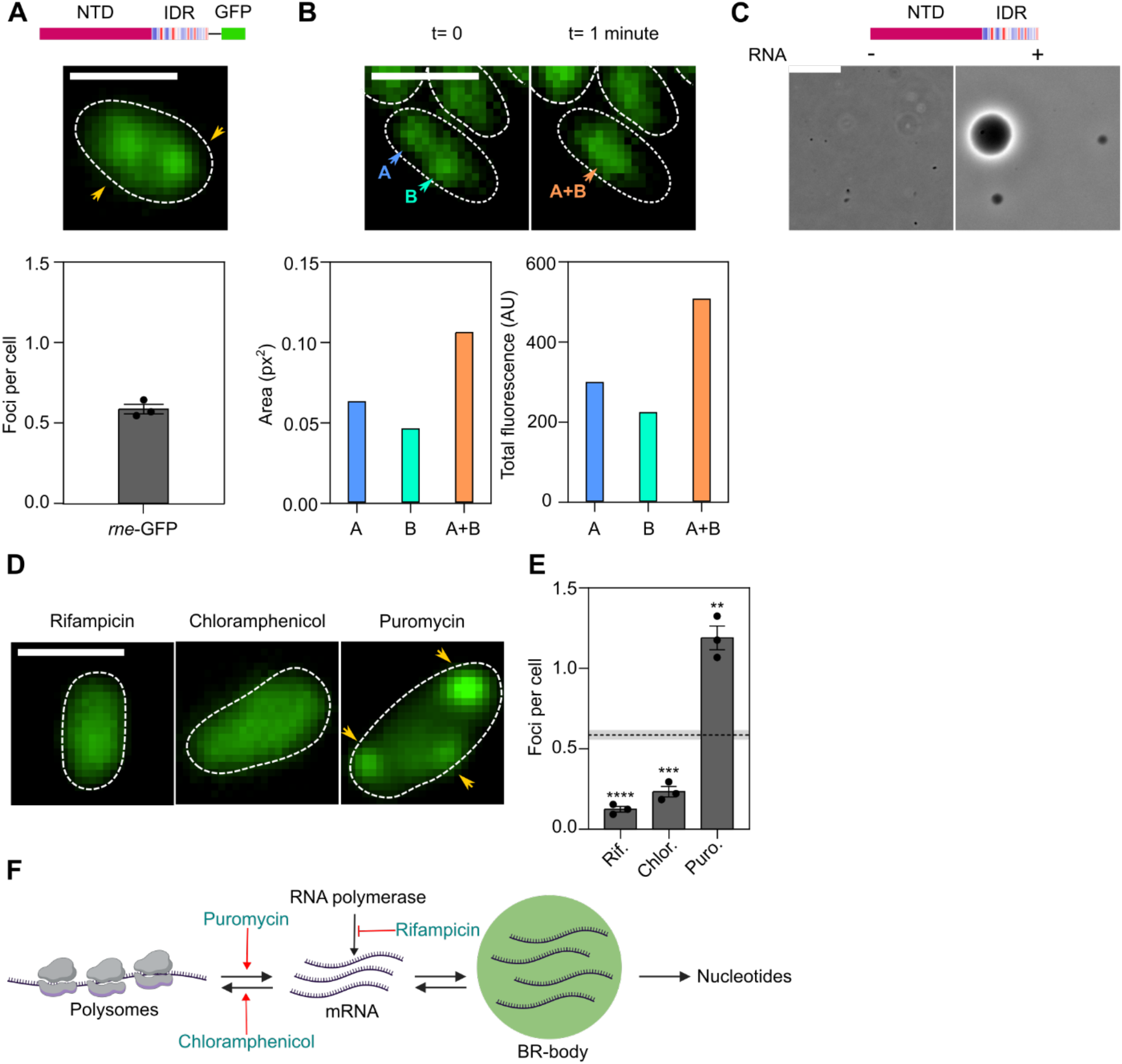
*B. ovis* RNase E forms RNA dependent condensates. **A)** Representative fluorescence microscopy image of a *B. ovis* cell expressing RNase E-GFP from the native *rne* locus (*rne*-GFP), with manual quantification of RNase E foci per cell from 401 cells across three biological replicates. Error bars represent the standard error of the mean (SEM). Scale bar, 1 μm. **B.)** Representative time-lapse fluorescence microscopy images showing the fusion of two RNase E-GFP foci over a 1-minute interval, with corresponding measurements of foci area and total fluorescence intensity within foci before and after fusion. Scale bar, 1 μm. **C)** *In vitro* condensate formation by purified full-length RNase E in the presence of RNA. Scale bar, 10 μm. **D)** Representative fluorescence microscopy images of *rne*-GFP cells following treatments with drugs that alter mRNA availability: rifampicin (100 μg/mL, 30 min) to inhibit transcription, chloramphenicol (100 μg/mL, 15 min) to inhibit translation and sequester mRNAs in polysomes, and puromycin (150 μg/mL, 30 min) to release mRNAs from polysomes. Scale bar, 1 μm. **E)** Manual quantification of RNase E foci per cell in drug-treated populations compared with untreated cells. The horizontal dashed line represents the average number of RNase E foci per cell in untreated cells shown in Fig. 2A. The following numbers of cells were analyzed across three biological replicates: rifampicin (n = 341), chloramphenicol (n = 327), and puromycin (n = 223). Error bars represent the standard error of the mean (SEM). p-values were calculated using a two-tailed *t* test with unequal variance (**, p < 0.01; ***, p < 0.001; ****, p < 0.0001). **F)** Schematic summarizing how rifampicin, chloramphenicol, and puromycin perturb cellular mRNA availability and the consequences for BR-body assembly.

To further explore the role of RNA in BR-body assembly *in vivo*, we imaged *B. ovis* BR-bodies upon treatment with various transcription and translation inhibitors. Poorly translated mRNAs have been reported to drive BR-body condensation in α-proteobacteria (12, 24), and we asked whether this was similarly the case in *B. ovis*. Treatment with rifampicin, which depletes free mRNA, or chloramphenicol, which stabilizes mRNA in polysomes, caused rapid dissolution of RNase E-GFP foci and redistribution of fluorescence throughout the cytoplasm (Fig. 2D-E). These results indicated that free untranslated mRNAs would promote BR-body assembly (Fig. 2F). Indeed, puromycin treatment, which releases nascent peptides from ribosomes and increases the pool of untranslated mRNA, enhanced BR-body formation (Fig. 2D). Quantification of foci per cell confirmed these trends across the population (Fig. 2E). To corroborate these results with an independent assay of RNase E condensation, we performed dual-color imaging of RNase E-GFP alongside constitutively expressed cytoplasmic dsRed (Fig. S1). Because *B. ovis* cells are small and spheroid, fluorescent signal appears brighter at the cell center due to increased path length. dsRed is known to be diffuse in the cytoplasm and is thus a useful reference for distinguishing true RNase E-GFP focus formation from this optical artifact. When RNase E-GFP condenses into foci its signal is expected to be poorly correlated with diffuse dsRed, whereas a diffuse RNase E-GFP distribution should produce high correlation with dsRed. Consistent with the foci measurements above, RNase E-GFP signal was poorly correlated with dsRed in untreated and puromycin-treated cells, and strongly correlated following rifampicin or chloramphenicol treatment (Fig. S1), supporting the conclusion that *B. ovis* BR-body assembly is stimulated by untranslated mRNA (12, 24)

### The C-terminal IDR of *B. ovis* RNase E is necessary and sufficient for BR-body phase separation

In other α-proteobacteria, the C-terminal IDR is required for BR-body assembly (12, 24), and we asked whether the same is true in *B. ovis*. A mutant lacking the RNase E IDR fused with a C-terminal monomeric superfolder GFP (*rne*(ΔIDR)-GFP) failed to form discrete RNase E-GFP foci and instead showed diffuse cytoplasmic signal (Fig. 3A, S2A-C), demonstrating that the IDR is necessary for BR-body formation in cells. Overall fluorescence from the RNase E(ΔIDR)-GFP fusion increased 2.2-fold relative to full-length RNase E-GFP (Fig. S2D-E), indicating that BR-body assembly is required for negative autoregulation of RNase E expression; negative autoregulation of RNase E has been previously reported in other bacterial genera (44–47). These data indicate that the *B. ovis* RNase E IDR contributes both to BR-body assembly and to proper autoregulatory control of RNase E protein levels.

**Figure 3.**
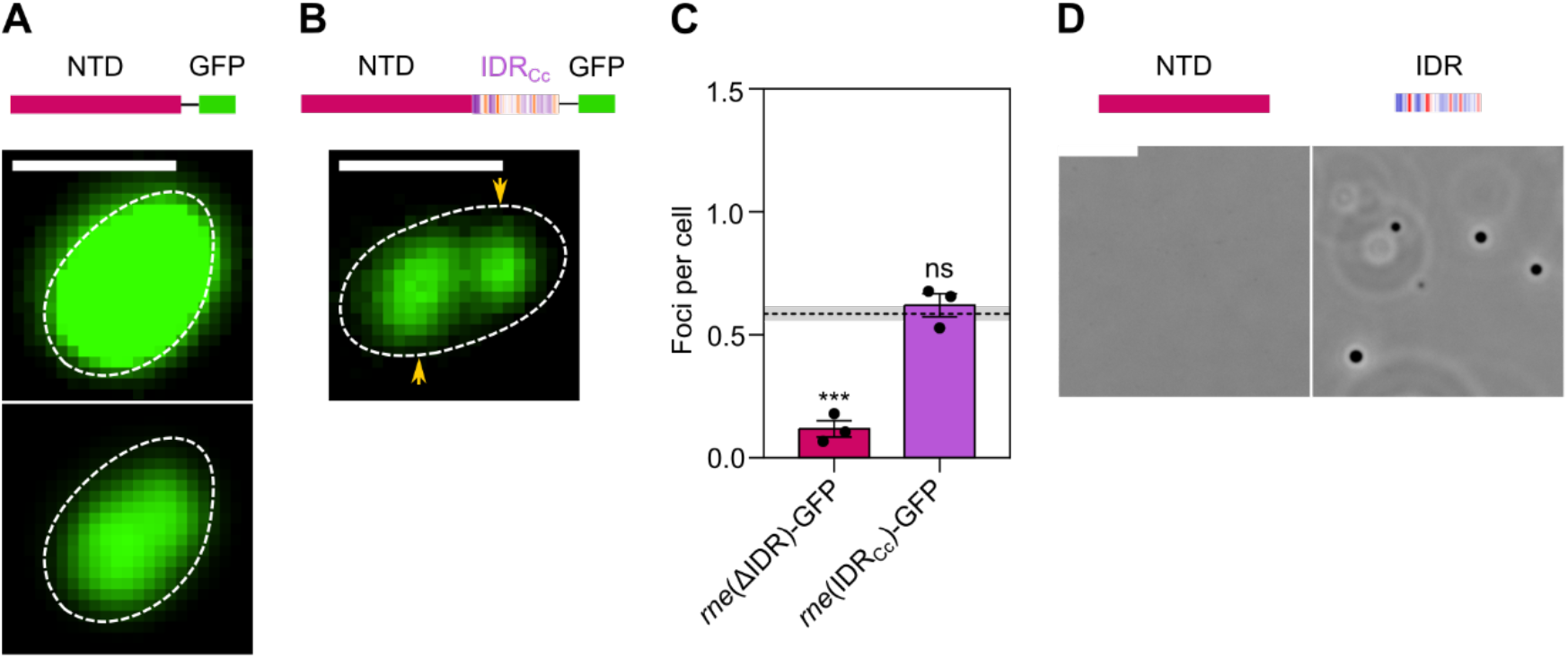
*B. ovis* RNase E C-terminal IDR is necessary and sufficient for BR-body phase separation. **A)** Representative fluorescence microscopy image of a *B. ovis* cell expressing RNase E(ΔIDR)-GFP from the native *rne* locus (*rne*(ΔIDR)-GFP). The top panel shows *rne*(ΔIDR)-GFP using the same display scale as *rne*-GFP in Fig. 2A. In the bottom panel, the display levels for *rne*(ΔIDR)-GFP were adjusted independently to improve image clarity. Scale bar, 1 μm. **B)** Representative fluorescence microscopy image of a *B. ovis* cell expressing a chimeric RNase E in which the C-terminal IDR is replaced with the IDR from *C. crescentus* RNase E and fused to a C-terminal GFP tag (*rne*(IDR_Cc_)-GFP). Scale bar, 1 μm. **C)** Manual quantification of foci per cell in *rne*(ΔIDR)-GFP and *rne*(IDR_Cc_)-GFP strains from three biological replicates (*n* = 484 and 195 cells, respectively), compared with *rne*-GFP cells from Fig. 2A. The horizontal dashed line represents the average number of foci per cell in untreated *rne*-GFP cells shown in Fig. 2A. Error bars represent the standard error of the mean (SEM). p-values were calculated using a two-tailed *t* test with unequal variance (***, p < 0.001; ns, not significant). **D)** *In vitro* condensate formation by purified *B. ovis* RNase EΔIDR and the isolated RNase E IDR in the presence of RNA. Scale bar, 10 μm.

To test whether a heterologous IDR is sufficient to drive phase separation of RNase E, we replaced the *B. ovis* IDR with the highly divergent *C. crescentus* IDR, which shares only 30% overall amino acid identity with the *B. ovis* IDR, and fused it with a C-terminal GFP (*rne*(IDR_Cc_)-GFP) (Fig. S3). The *C. crescentus* IDR is known to drive phase separation of its RNase E protein (12). The chimeric RNase E(IDR_Cc_) formed BR-bodies (Fig. 3B-C; Fig. S2A-C), demonstrating that a heterologous IDR can drive RNase E condensation in *B. ovis* cells (Fig. 3B-C). The RNase E(IDR_Cc_) chimera protein is expressed (Fig. S2D-F), and does not show the large autoregulation defect observed in the *B. ovis rne*(ΔIDR)-GFP mutant.

Finally, we tested whether the *B. ovis* IDR is sufficient for phase separation *in vitro* by examining purified IDR and catalytic domain fragments. Although the *B. ovis* IDR has very low primary structure identity with other α-proteobacterial IDRs, it preserves the characteristic alternating blocks of positively and negatively charged residues previously shown to underlie RNase E liquid-liquid phase separation (12, 15) (Fig. 1D). Under conditions where full-length RNase E formed droplets in the presence of RNA (Fig. 2C), the isolated catalytic domain failed to phase separate and remained uniformly distributed in solution, while the isolated IDR formed spherical condensates with RNA (Fig. 3D). Together, these results demonstrate that the C-terminal IDR of *B. ovis* RNase E is necessary for BR-body assembly in cells, sufficient to form phase separated droplets with RNA *in vitro*, and that the highly divergent *C. crescentus* IDR supports RNase E condensation in *B. ovis*. These results further provide evidence that IDR-dependent BR-body assembly contributes to negative autoregulation of RNase E levels.

### RNase E BR-bodies protect *B. ovis* against oxidative, membrane, and cell wall stresses

Deletion of the C-terminal IDR domain of RNase E is tolerated during exponential growth under standard laboratory conditions in several genera (12, 24, 25, 36). Consistent with this, our analysis of published Tn-Himar sequencing datasets from *B. ovis* (48) and *B. abortus* (49, 50) revealed that transposon insertions are readily recovered within the IDR-encoding region of RNase E, while insertions within the conserved N-terminal catalytic nuclease domain are entirely absent (Fig. 4A). These data indicate that the IDR (and by extension BR-body formation) is dispensable for viability under standard unstressed culture conditions, whereas the catalytic nuclease core is essential.

**Figure 4.**
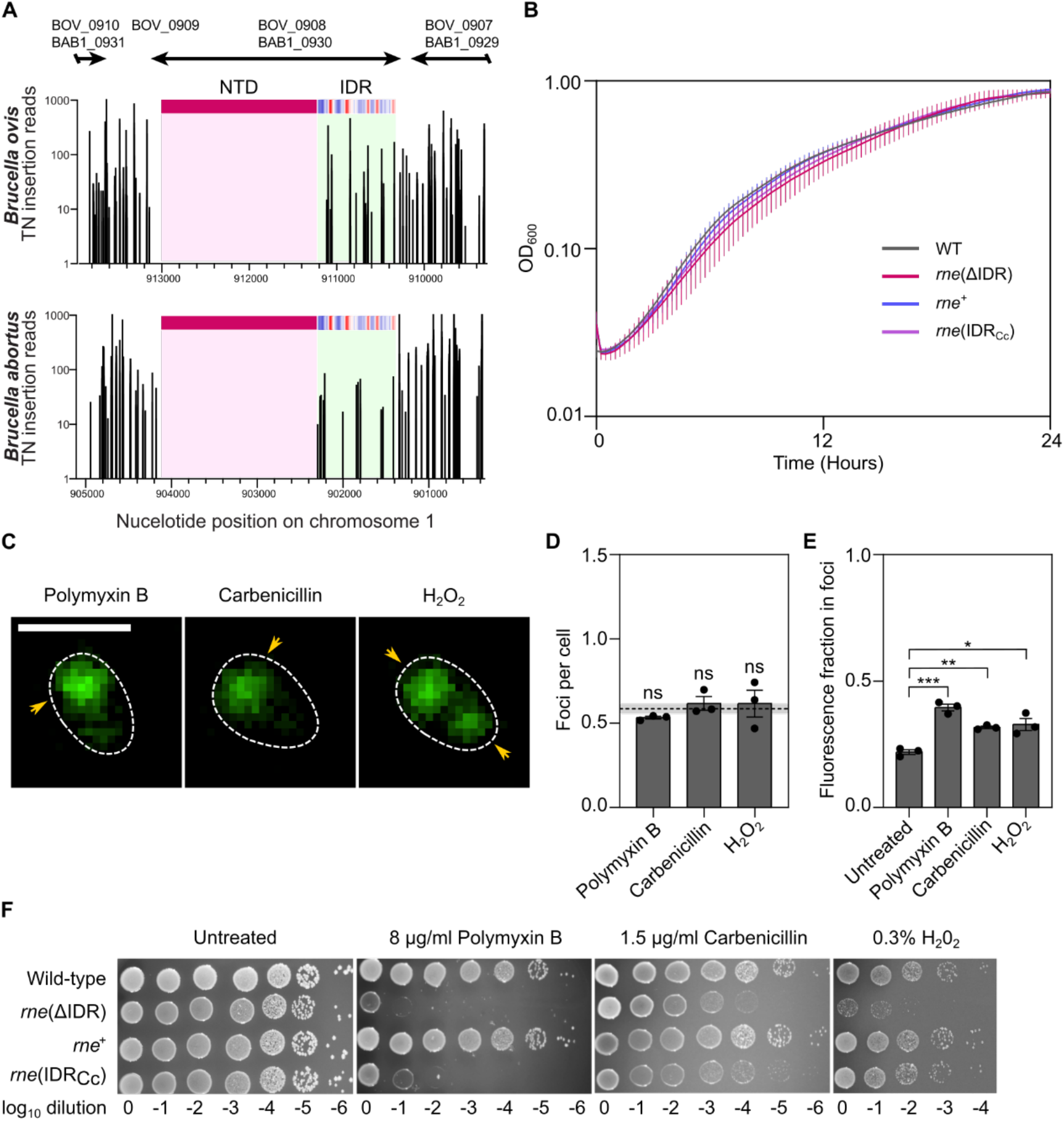
*B. ovis* BR-bodies contribute to stress tolerance. **A)** Transposon insertion profiles across the *rne* locus on chromosome 1 in *B. ovis* and *B. abortus*. Locus annotations for *B. ovis* (BOV_0908) and *B. abortus* (BAB1_0930) are indicated above the plot. The RNase E N-terminal catalytic nuclease domain and the C-terminal IDR are highlighted (pink and light green, respectively). Per position Tn insertion read counts were calculated as previously described for these Tn-seq datasets (48, 50, 70). **B)** Growth of the wild-type strain, *rne*(ΔIDR), the genetically restored *rne*^+^ strain, and the *rne*(IDR_Cc_) chimeric strain in broth culture under standard (unstressed) conditions, monitored by OD_600_. **C.)** Representative fluorescence microscopy images of *rne*-GFP cells following treatment with polymyxin B (8 µg/mL, 30 min), H_2_O_2_ (0.3%, 30 min), or carbenicillin (1.3 µg/mL, 30 min). Scale bar, 1 μm. **D.)** Manual quantification of foci per cell following treatment with polymyxin B (8 µg/mL, 30 min), H_2_O_2_ (0.3%, 30 min), or carbenicillin (1.3 µg/mL, 30 min), compared with untreated cells from Fig. 2A. The horizontal dashed line represents the average number of foci per cell in untreated *rne*-GFP cells shown in Fig. 2A. Cells analyzed from three biological replicates were as follows: untreated (n = 401), polymyxin B (n = 296), H_2_O_2_ (n = 164), and carbenicillin (n = 238). Error bars represent the standard error of the mean (SEM). p-values were calculated using a two-tailed *t* test with unequal variance (ns, not significant). **E.)** Fraction of RNase E–GFP fluorescence localized to foci following exposure to polymyxin B (8 µg/mL, 30 min), H_2_O_2_ (0.3%, 30 min), or carbenicillin (1.3 µg/mL, 30 min), compared with untreated cells. Thirty cells per condition from three biological replicates were analyzed. Error bars indicate the standard error of the mean (SEM). p-values were calculated using a two-tailed *t* test with unequal variance (***, p < 0.001; **, p < 0.01; *, p < 0.05). **F.)** Spot dilution assays of wild type, *rne*(ΔIDR), *rne*⁺, and the *rne*(IDR_Cc_) chimera for stress sensitivity. Strains grown on TSA blood agar were resuspended, normalized to OD_600_ = 0.3, serially diluted 10-fold, and spotted (5 µL) onto TSA blood agar plates lacking stressor or supplemented with polymyxin B (8 µg/mL), carbenicillin (1.5 µg/mL), or H_2_O_2_. Plates were incubated at 37°C and imaged after 72-96 hours. Dilution plating experiments were repeated at least three times (n=3) in technical triplicate, and one representative experiment is shown.

To directly test whether BR-bodies impact *B. ovis* fitness, we compared the BR-body-deficient *rne*(ΔIDR) mutant to isogenic strains carrying either the restored native IDR (*rne*^+^) or the chimeric allele in which the *B. ovis* IDR was replaced with the *C. crescentus* IDR (*rne*(IDR_Cc_). Given that the RNase E IDR not only drives phase separation but also scaffolds recruitment of degradosome protein and RNA clients (11, 51), these two functions must be separated to understand which aspects of IDR function support *B. ovis* fitness. The *rne*(IDR_Cc_) chimera therefore served as a separation-of-function allele to ask whether phase separation *per se* is sufficient for RNase E function in *B. ovis*. Although the composition of the *B. ovis* RNA degradosome remains undefined, the *C. crescentus* RNase E IDR binding sites for degradosome clients are poorly conserved in *B. ovis*, with aconitase, RNase D, and PNPase interaction motifs sharing only 39%, 40%, and 60% identity, respectively (Fig. S3). The *rne*(IDR_Cc_) chimera therefore retains phase separation capacity but is expected to be impaired in recruitment of *B. ovis*-specific degradosome components.

All three strains exhibited similar growth kinetics in exponential-phase broth culture and reached similar final optical densities compared to wild-type *B. ovis* (Fig. 4B), suggesting that the deletion of the IDR domain of RNase E and loss of BR-body formation does not measurably impair *B. ovis* growth under standard (unstressed) broth culture conditions.

BR-bodies can be dynamically remodeled by environmental stress and become more prominent under conditions that inhibit translation or damage macromolecules (12, 26). To test whether *B. ovis* BR-bodies are stress responsive, we imaged RNase E-GFP condensates in cells exposed to oxidative stress (H_2_O_2_), the cationic antimicrobial peptide polymyxin B, or the β-lactam antibiotic carbenicillin. Under all three conditions, the number of RNase E foci or the total fluorescence within cells did not change significantly, but fluorescence fraction in foci intensity increased (Fig. 4C-E), indicating that BR-body dynamics are altered upon stress; this suggested a role for BR-bodies in stress responses. We therefore tested whether BR-body formation contributes to survival under these stress conditions.

The *rne*(ΔIDR), *rne*^+^, *rne*(IDR_Cc_) chimera, and the wild-type strains were spotted onto regular agar plates, agar containing polymyxin B or carbenicillin, or were challenged with H_2_O_2_, and colony growth was assessed. Colony morphology and plating efficiency on untreated control conditions were comparable across all strains, confirming that the RNase E IDR is dispensable for growth on standard solid medium (Fig. 4F). In contrast, the *rne*(ΔIDR) mutant was markedly more sensitive to all three stressors than wild-type, exhibiting smaller colonies and reduced recovery at higher stress concentrations (Fig. 4F). Restoring the native *B. ovis* IDR fully rescued these defects, demonstrating that the RNase E IDR is required for resistance to oxidative, membrane, and cell wall stress. The *rne*(IDR_Cc_) chimera variably rescued stress sensitivity: oxidative stress resistance was restored to near wild-type levels, while resistance to polymyxin B and carbenicillin was only marginally improved (Fig. 4F). These results indicate that RNase E phase separation is sufficient to promote oxidative stress tolerance, but that full resistance to antimicrobial peptide and cell wall stress additionally requires lineage-specific IDR functions, such as recruitment of *B. ovis*-specific degradosome components. We conclude that *B. ovis* BR-bodies are dispensable for growth under standard unstressed culture conditions but are important for mounting an effective response to infection-relevant stresses *in vitro*.

### *Brucella* BR-bodies primarily promote mRNA decay

To test whether BR-bodies influence RNA decay in *B. ovis*, we used Rif-seq to measure transcript half-lives in exponential-phase cultures, comparing wild-type cells to the BR-body-deficient *rne*(ΔIDR) mutant (Fig. 5; Table S1). BR-bodies had a predominantly destabilizing effect on the *B. ovis* transcriptome: among transcripts whose decay rates were significantly altered in the *rne*(ΔIDR) mutant, ∼74% displayed increased half-lives, while ∼26% were destabilized (Fig. 5A; Table S1). These data indicate that under the conditions tested, BR-bodies primarily promote mRNA turnover rather than stabilization, supporting a conserved role for BR-bodies as hubs that accelerate mRNA decay across α-proteobacteria.

**Figure 5.**
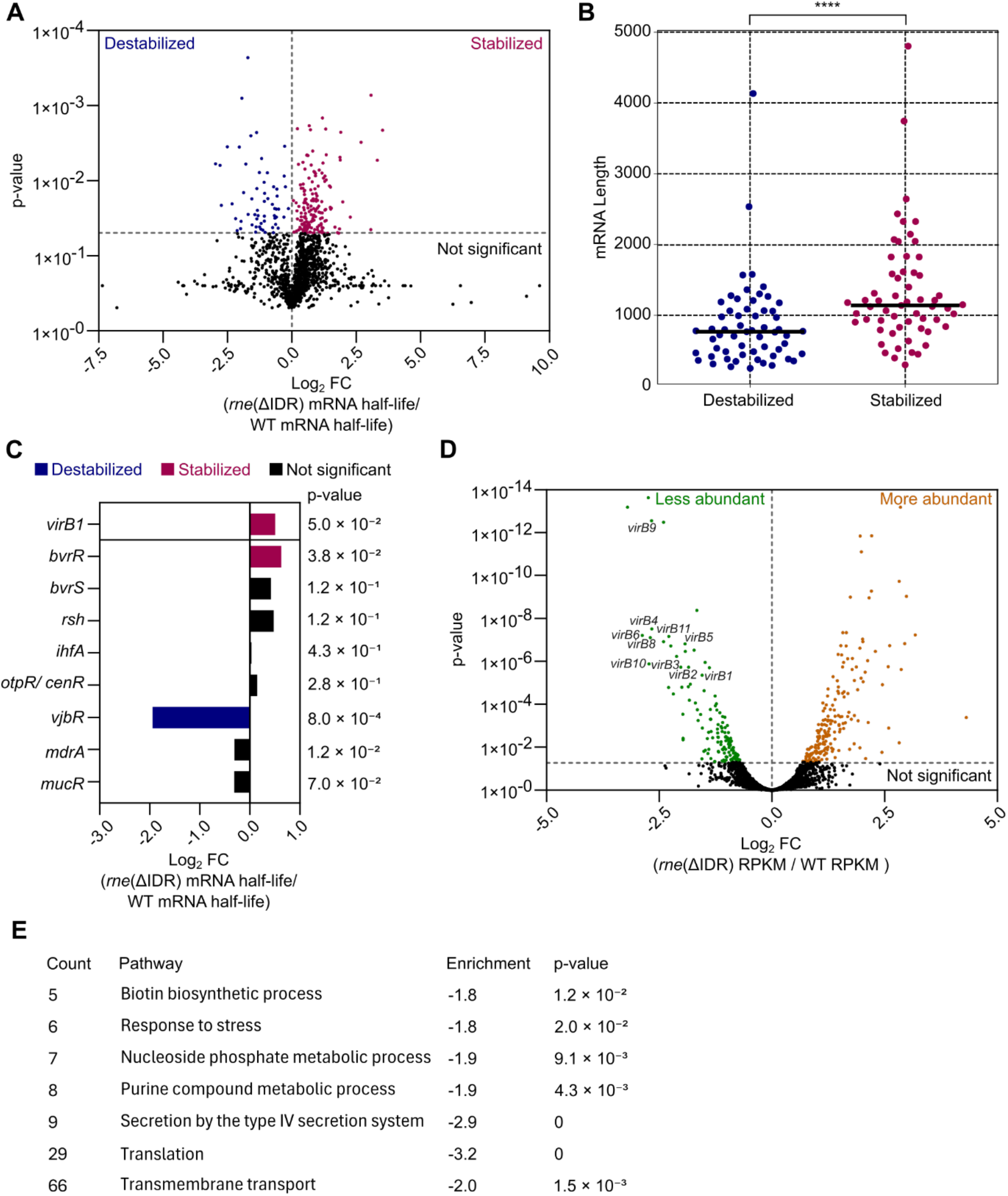
Loss of BR-bodies broadly remodels mRNA stability, including the *virB* operon and its upstream regulators through direct and indirect control. **A**) Fold changes in mRNA half-lives for 1,524 transcripts measured by Rif-seq in wild-type or *rne*(ΔIDR) strains. Significance was assessed at a nominal p < 0.05 using a one-tailed *t* test with unequal variance. Each point represents a single trasncript and the points are colored by significance class (stabilized, destabilized, or not significant as indicated on the plot) **B)** Comparison of transcript lengths for mRNAs classified as significantly stabilized or significantly destabilized in the *rne*(ΔIDR) strain based on Rif-seq half-life measurements. p-values were calculated using a one tailed t test with unequal variance. (****, p < 0.0001). **C)** Differential stabilization of regulators of the *Brucella virB* operon and *virB1* in the *rne*(ΔIDR) mutant. Bars are colored by significance class (stabilized, destabilized, or not significant as indicated on the plot) and the p-values were calculated using a one tailed *t* test with unequal variance. **D)** Volcano plot of differential gene expression comparing *rne*(ΔIDR) to wild type (WT) using DESeq2. The x-axis shows log_2_ fold change (*rne*(ΔIDR)/WT), and the y-axis shows the corresponding p-values plotted on a logarithmic scale. Each point represents a single transcript and the points are colored by significance class (more abundant, less abundant, or not significant as indicated on the plot; Benjamini–Hochberg–adjusted p < 0.05), and selected T4SS (*virB*) operon genes are labeled. **E)** Gene set enrichment analysis (GSEA) of GO terms using OmicsBox. All genes were ranked by log_2_ fold change and used as input for enrichment testing. GO terms are grouped by Biological Process (BP), Cellular Component (CC), and Molecular Function (MF) and only the Biological Process group is shown here. Because *virB* operon genes have limited GO annotation, GO:0043684 (type IV secretion system) and GO:0044097 (secretion by type IV secretion system) were manually assigned to *virB* locus tags as described in Methods. p-values were adjusted for multiple testing using the Benjamini-Hochberg procedure to control the false discovery rate.

We next asked whether BR-bodies act preferentially on specific classes of transcripts. In *C. crescentus*, BR-body-enriched RNAs tend to be long and poorly translated, and these transcripts are selectively stabilized upon IDR deletion (13). We observed a similar pattern in *B. ovis*; mRNAs with increased half-lives in the *rne*(ΔIDR) strain were significantly longer than destabilized transcripts (Fig. 5B). This result provides evidence that BR-bodies in *Brucella* also specialize in degrading long mRNAs, supporting a model in which RNase E BR-bodies preferentially engage long mRNAs, likely due to increased multivalency which can promote phase-separation.

### BR-bodies directly and indirectly impact translation, transport, and virulence processes

The type IV secretion system (T4SS) is a critical virulence determinant in *B. ovis* (52), with its structural components encoded by the long, polycistronic *virB* operon. Since BR-bodies preferentially promote the decay of long mRNAs, we examined the stability of *virB* mRNA in the *rne*(ΔIDR) mutant relative to wild-type. Among the *virB* transcripts, only *virB1* was detected with sufficient coverage for half-life analysis in both strains, and its mRNA was significantly stabilized in the *rne*(ΔIDR) BR-body-deficient mutant, consistent with its impaired mRNA decay (Fig. 5C). As presented below, however, *virB1* and the downstream T4SS genes were reduced in abundance in the mutant at steady-state (Fig. 5D), indicating that the net expression of the *virB* operon decreases upon loss of BR-bodies despite this local stabilization. The expression of the *virB* operon is controlled by a complex regulatory network involving multiple upstream transcriptional regulators; many of these regulators also exhibited altered mRNA stability in the *rne*(ΔIDR) mutant (Fig. 5C), suggesting that disruption of BR-body formation influences T4SS expression through both direct stabilization of *virB* transcripts and through effects on expression of regulators that control *virB* gene expression.

To complement the Rif-seq half-life measurements and capture the cumulative effects of altered RNA metabolism on steady-state mRNA levels, we performed RNA-seq on wild-type and *rne*(ΔIDR) strains. Gene Ontology (GO) enrichment analysis was conducted on transcripts whose steady-state levels were significantly increased or decreased in the *rne*(ΔIDR) mutant relative to wild type (Fig. 5D). Transcripts that accumulated in the *rne*(ΔIDR) mutant were strongly enriched for membrane- and transport-related functions. Specifically, the accumulated set included numerous ABC transporter operons, tripartite ATP-independent periplasmic (TRAP) transporters, and branched-chain amino acid and sulfate uptake systems. We also observed over-representation of peptidoglycan L,D-transpeptidase activity and peptidoglycan-protein cross-linking, implicating cell wall remodeling enzymes among the transcripts that normally undergo rapid BR-body-dependent turnover (Table S2 and S3). We conclude that under standard cultivation conditions, the RNA degradosome post-transcriptionally limits expression of select envelope transport and nutrient uptake systems, as well as envelope-modifying enzymes that impact cell wall architecture.

In contrast, transcripts with reduced abundance in the *rne*(ΔIDR) mutant were enriched for core biosynthetic and virulence-associated functions (Table S3). GO terms related to translation, including structural constituents of the ribosome and translation factors, were significantly over-represented among mRNAs with reduced steady-state levels, suggesting that BR-bodies contribute to the maintenance of transcripts encoding the core translation machinery. Transcripts encoding biotin biosynthetic enzymes and components of the VirB type IV secretion system (T4SS), a critical virulence determinant across *Brucella* spp. (53–56), were also reduced at steady state in the *rne*(ΔIDR) mutant (Table S3). A possible mechanism underlying reduced *virB* transcript levels involves *vjbR*, which encodes a major transcriptional activator of the T4SS (57, 58). *vjbR* had lower steady-state levels and its mRNA was strongly destabilized in the *rne*(ΔIDR) mutant (Fig. 5C), providing evidence that BR-bodies normally protect *vjbR* transcripts from decay.

To complement the GO overrepresentation analyses above, we also performed Gene Set Enrichment Analysis (GSEA), which ranks all genes by their steady-state log_2_ fold change (*rne*(ΔIDR) versus wild-type) to identify gene sets with coordinated expression changes without imposing a significance threshold (Table S4). Consistent with the GO enrichment results, gene sets associated with biotin biosynthesis and translation were enriched toward the bottom of the ranked list, whereas gene sets related to transmembrane transport, including ABC transporters, were enriched toward the top of the ranked list. Most notably, GSEA identified significant negative enrichment of the type IV secretion system gene set, indicating that T4SS-associated genes are coordinately shifted toward lower expression in the *rne*(ΔIDR) mutant (Fig. 5E, S5). Together, these complementary genome-scale analyses support our conclusion that BR-bodies control the homeostasis of key biosynthetic and virulence transcripts, either through direct stabilization, degradation, or through higher-order regulatory mechanisms that control the levels upstream transcriptional and post-transcriptional regulators.

To gain higher-resolution insight into BR-body-dependent regulation of the large *virB* operon, we took advantage of the end-enriched (REND-seq) design of our Rif-seq libraries (39), which maps transcript 5ʹ and 3ʹ ends at single-nucleotide resolution in wild-type and *rne*(ΔIDR) cells. Across the operon, *virB*-mapped reads decreased more rapidly in the *rne*(ΔIDR) mutant, consistent with accelerated decay of *virB* transcripts under standard cultivation conditions (Fig. 6). Mapping of 5ʹ ends further revealed two prominent putative processing sites near the 5ʹ ends of *virB2* and *virB5*, each positioned adjacent to an antisense RNA (Fig. 6). This positional relationship suggests that antisense transcription (or the antisense transcripts themselves) may regulate RNase E-dependent cleavage of the *virB* operon at specific sites, suggesting a possible antisense-associated mechanism for site-specific *virB* processing.

**Figure 6.**
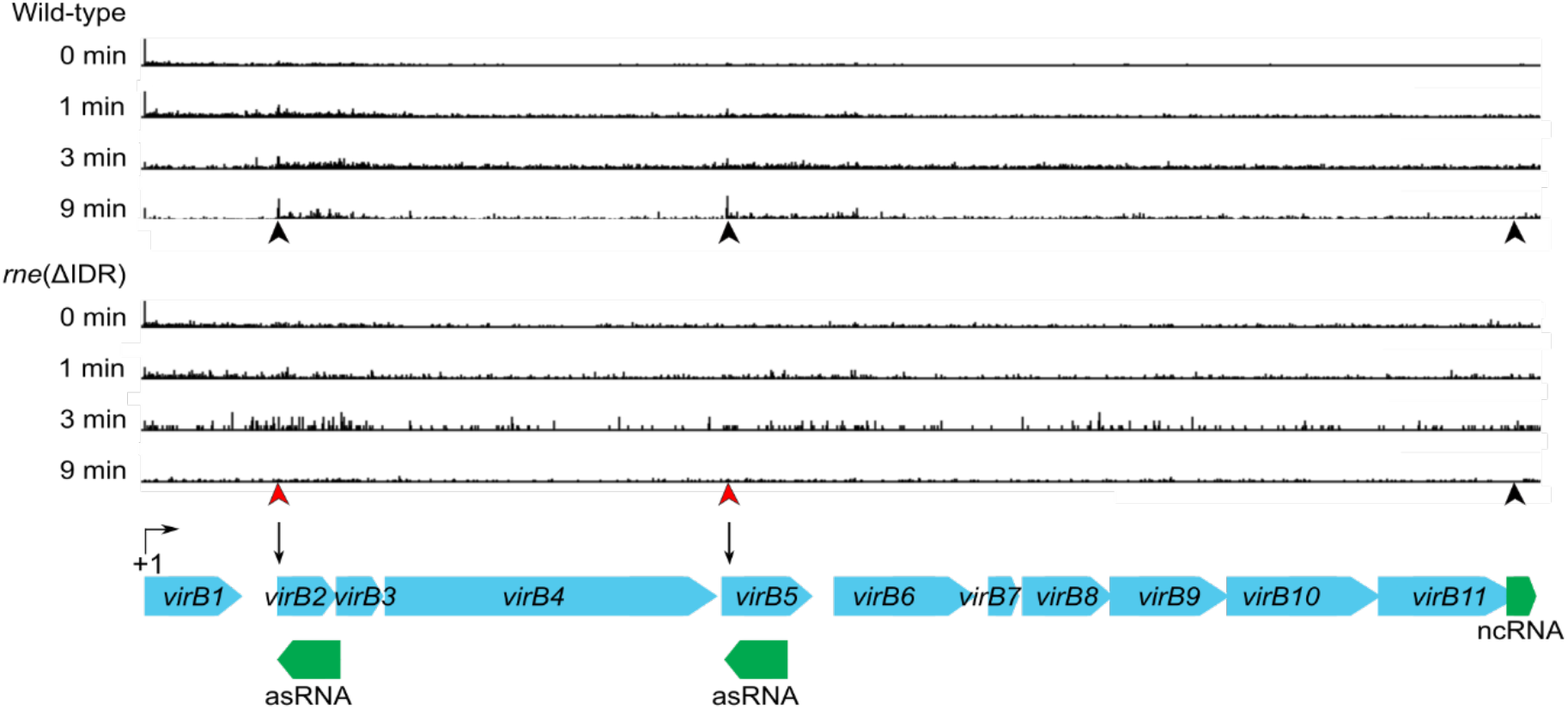
RNase E IDR influences potential processing within the *Brucella virB* (T4SS) operon. Genome browser view of 5’ REND-seq RNA end profiles across the complete *virB* locus in wild-type *B. ovis* (top) and the RNase E IDR deletion strain (*rne*(ΔIDR); bottom) collected at 0, 1, 3, and 9 minutes after transcriptional shutoff by rifampicin (timepoints indicated at left). Tracks show mapped 5’ signal across the operon (vertical black lines). Tracks show a representative biological replicate per genotype (two biological replicates were analyzed per genotype), and the vertical axis reflects 5’-end read depth (normalization as described in Methods). Red arrows denote prominent RNA end positions interpreted as putative processing sites, including sites near the 5ʹ regions of *virB2* and *virB5*. Gene annotations for the *virB* operon (*virB1*-*virB11*) are shown in cyan; annotated antisense RNAs detected by RNA-sequencing adjacent to the *virB2* and *virB5* regions (asRNA1 and asRNA2) are shown in green, along with a stable non-coding RNA (ncRNA) feature downstream of *virB11*.

### BR-bodies promote intracellular fitness in mammalian macrophages

Given the central role of BR-bodies in *B. ovis* RNA metabolism, stress resistance and virulence gene expression, we next asked whether these condensates support fitness in the intracellular niche. We quantified the fitness of the *B. ovis rne*(ΔIDR) mutant in THP-1 macrophage-like cells by monitoring CFUs over time (Fig. 7). Although the *rne*(ΔIDR) mutant entered macrophages at levels comparable to wild type, its ability to survive and/or replicate intracellularly was severely impaired. At both 24 and 48 hours post-infection, CFU counts for *rne*(ΔIDR) were significantly reduced relative to the wild-type strain and the genetically restored control (*rne*^+^), indicating that BR-body formation promotes fitness within the intracellular niche.

**Figure 7.**
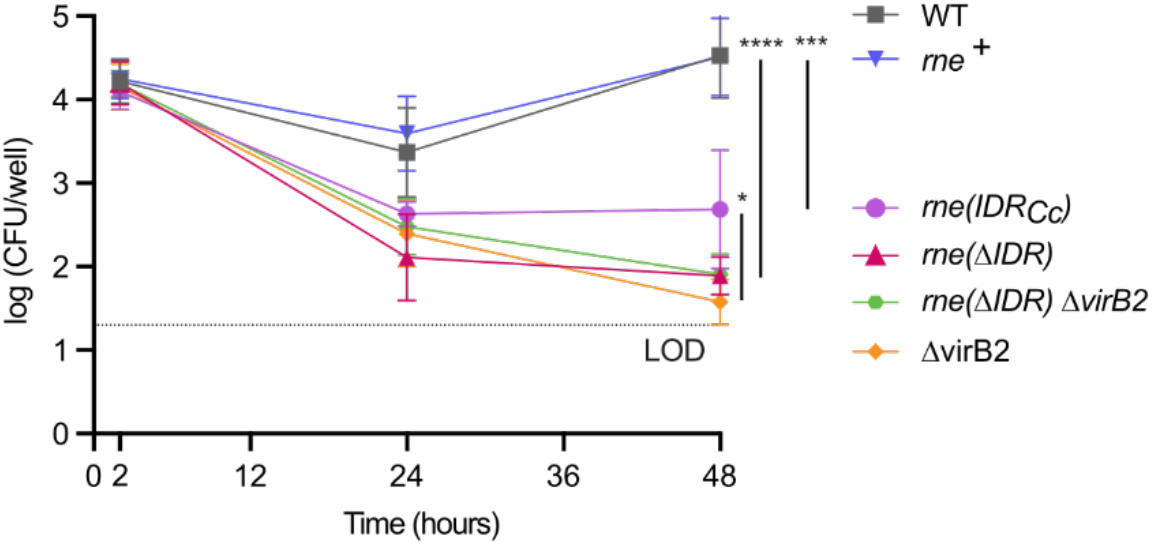
BR-body phase separation and client recruitment support *B. ovis* intracellular fitness in THP-1 macrophages. THP-1 macrophage-like cells were infected with wild-type *B. ovis* (WT, black squares), an RNase E IDR deletion mutant (*rne*(ΔIDR), blue triangles), the RNase E restored strain (*rne^+^,* red inverted triangles), the RNase E chimera (*rne*(IDR_Cc_)) (purple circles), a T4SS-defective Δ*virB2* mutant (orange diamonds), or the *rne*(ΔIDR) Δ*virB2* double mutant (green circles). Intracellular bacteria were quantified at 0, 24, and 48 hours post-infection by enumerating colony-forming units (CFU) per well and are plotted on a log_10_ scale. Data points represent mean CFU ± SD of three biological replicates. Limit of detection (LOD) for the assay is shown as a horizontal dotted line. CFUs at the 48-hour timepoint were compared using a one-way ANOVA followed by Tukey’s post hoc test. p-values for the comparisons between strains are shown (*, p < 0.05; ***, p < 0.001; ****, p < 0.0001).

Given that *rne*(ΔIDR) also exhibits strong dysregulation of *virB* operon transcripts (Fig. 5; Table S2), we hypothesized that its intracellular defect is caused, at least in part, by impaired type IV secretion system (T4SS) function. To test this model, we constructed a Δ*virB2* mutant, which lacks a functional T4SS, as well as a *rne*(ΔIDR) Δ*virB2* double mutant. The double mutant showed no additional attenuation relative to either single mutant (Fig. 7), consistent with the T4SS pathway contributing substantially to the intracellular fitness defect of *rne*(ΔIDR). In contrast, Δ*virB2* did not display the polymyxin B or carbenicillin sensitivity seen in *rne*(ΔIDR) (Fig. S4), indicating that T4SS perturbation alone does not explain the general *in vitro* stress sensitivity associated with loss of BR-bodies.

Finally, we leveraged the *rne*(IDR_Cc_) chimera strain to test whether RNase E condensation was sufficient to support intracellular fitness. The *rne*(IDR_Cc_) strain only partially rescued the *rne*(ΔIDR) fitness defect in macrophages (Fig. 7), indicating that RNase E phase separation alone is insufficient to support full *B. ovis* fitness in the intracellular niche. Collectively, these data support a model in which the *Brucella* RNase E IDR supports *B. ovis* in the macrophage by promoting BR-body assembly and regulating infection-relevant degradosome clients including the *virB* operon mRNA. The failure of the heterologous *C. crescentus* IDR to fully rescue the intracellular fitness defect of *rne*(ΔIDR) shows that the native *Brucella* RNase E IDR is specifically required, likely because it couples condensate formation to recruitment of infection-relevant RNA degradosome clients.

## Discussion

### Role of bacterial condensates in host-colonization

Biomolecular condensates have emerged as important regulators of gene expression and stress adaptation across all domains of life, and growing evidence now links these structures to fitness during host colonization. Here we show that BR-bodies in the intracellular animal pathogen *B. ovis* play important roles in global mRNA decay, stress adaptation, and virulence gene regulation. These results extend previously reported attenuation of a *B. abortus* RNase E IDR truncation mutant in a mouse infection model (36), implicating BR-body-dependent RNA regulation as a conserved determinant of *Brucella* pathogenesis. More broadly, Tn-seq studies in a number of Gram-negative species have found that disrupting the BR-body-forming IDR of RNase E reduces host colonization (27–33), suggesting that support of fitness host contexts may be a widely shared function of BR-bodies. Beyond pathogens, BR-bodies in the plant symbiont *Sinorhizobium meliloti* likewise influence global mRNA decay, stress survival, and root colonization (24), demonstrating the importance of condensate-mediated RNA regulation in plant-bacteria interactions. Other bacterial biomolecular condensates have also been linked to post-transcriptional coordination of host colonization, including transcription termination factor Rho condensates that promote gut colonization in *Bacteroides* (59), and CsrA foci that support host cell colonization in enterohemorrhagic *E. coli* (60). Collectively, these observations suggest that biomolecular condensates are widespread organizers of stress response and virulence gene regulation in bacteria, and may represent promising targets for next-generation antimicrobial strategies.

### BR-bodies and type IV secretion system regulation

A *B. ovis* mutant that cannot produce BR-bodies (*rne*(ΔIDR)) exhibits dysregulation of the *virB* type IV secretion system (T4SS) (Figs. 5, 6). Secretion system mRNAs are frequently long and poorly translated, features that drive preferential partitioning of RNAs into BR-bodies across species (13, 24). Thus direct BR-body-mediated regulation of T4SS transcripts may be a general feature of pathogens that use these large secretion system operons to manipulate hosts. In *B. ovis*, BR-bodies appear to directly influence the stability of the *virB1* portion of the polycistronic *virB* message and are linked to distinct RNA processing events near the 5ʹ ends of *virB2* and *virB5*, potentially coordinated by antisense RNAs encoded within these CDSs (Figs. 5, 6). Whether these processing events promote or repress translation of the *virB2-11* CDSs remains to be determined. Notably, mRNA storage has been reported in *C. crescentus* BR-bodies during stress, with stored transcripts remaining translationally competent upon recovery (26). An important future direction will be to determine whether *virB* mRNAs are similarly stored within BR-bodies and released at specific stages of infection to coordinate *Brucella*-containing vacuole formation and intracellular replication.

BR-body disruption also dysregulated several upstream regulators of *virB*, including the direct activator VjbR (57, 58), the small regulatory RNA Bsr4 (61), the BvrR/BvrS two-component system (62), and Rsh (63) (Fig. 5C; Tables S1-S4). These changes likely compound the direct effects of BR-bodies on *virB* transcript levels and may have broader physiological impacts in the intracellular niche. Importantly, our experiments with the *rne*(IDR_Cc_) chimera demonstrate that formation of BR-body condensates alone is insufficient for full rescue of fitness during stress treatment or infection (Fig. 4 & 7). These results suggest that recruitment of *B. ovis*-specific RNA degradosome clients by the IDR is required to support fitness under select conditions, both *in vitro* and in host cells. The composition of the *Brucella* RNA degradosome remains undefined, though degradosome composition differs considerably across bacterial species. In *Rhizobium leguminosarum*, for example, an RNase E-containing complex associates with Hfq and RNA helicase(s) (64), while in *C. crescentus* the RNase E-associated degradosome includes PNPase, RNA helicases, aconitase, and RNase D (65–67). Thus, degradosome composition can vary even among α-proteobacteria. Defining the composition of the *B. ovis* RNA degradosome will therefore be an important step toward understanding how IDR-client interactions coordinate the post-transcriptional regulation of virulence and stress response mRNAs within BR-bodies. Altogether, our results establish BR-bodies as post-transcriptional hubs that link RNA metabolism to virulence and stress adaptation in *Brucella*. They further provide evidence that the specific degradosome-client interactions mediated by the RNase E IDR, not phase separation alone, are required for proper BR-body function during infection.

## Acknowledgements

This research was supported by a WSU Career Chair Award to J.M.S., as well as by the National Institute of General Medical Sciences of the NIH under award numbers R35GM124733 to J.M.S. and R35GM131762 to S.C. Support was also provided by the National Institute of Allergy and Infectious Diseases through awards F32AI174818 to M.A. and R01AI177619 to S.C.

## Supplementary Figures

**Figure S1:**
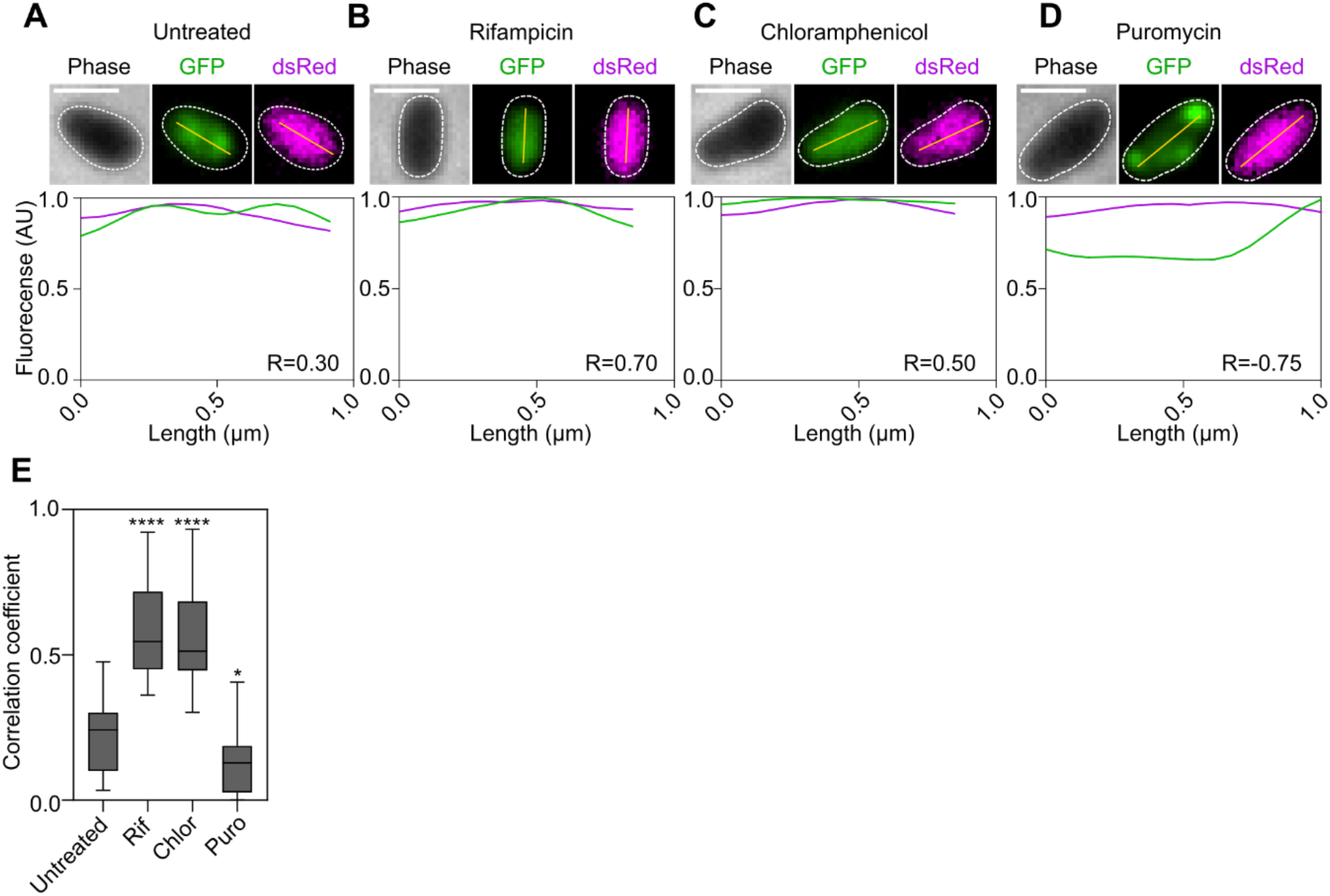
Line trace analysis of *rne*-GFP relative to cytoplasmic dsRed control. A-D) Phase-contrast and fluorescence microscopy images of dual-labeled *B. ovis rne*-GFP cells expressing full-length RNase E-GFP (green) together with a constitutive *glmS*::dsRed marker (magenta) in untreated **(A)**, rifampicin (100 µg/mL, 30 min) treated **(B)**, chloramphenicol (100 µg/mL, 15 min) treated **(C)**, or puromycin (150 µg/mL, 30 min) treated **(D)** cells. The yellow line indicates the transect used to generate the fluorescence intensity profiles shown below each image; line profiles were quantified in Fiji (ImageJ). Scale bars, 1 μm. (**E)** Colocalization between GFP and dsRed signals, expressed as Pearson correlation coefficients calculated from 60 cells across three biological replicates for untreated and drug treated conditions. Bars show mean values and error bars indicate standard deviation. (*, p< 0.05; ****, p < 0.0001, two-tailed *t* test with unequal variance).

**Figure S2:**
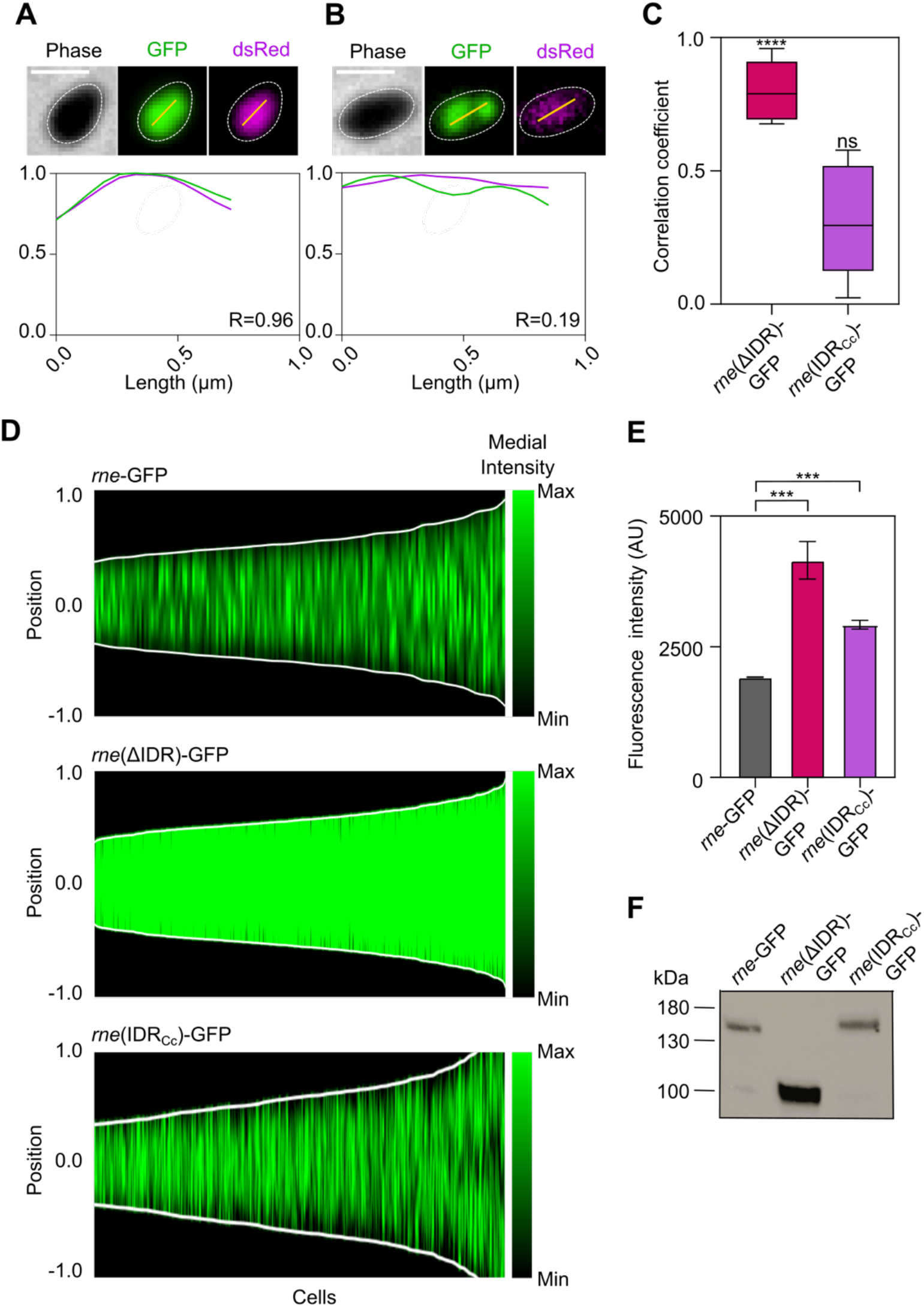
*rne*(ΔIDR)-GFP and *rne*(IDR_Cc_)-GFP expression and localization. **A)** Phase-contrast and fluorescence images of dual-labeled *rne*(ΔIDR)-GFP cells expressing RNase EΔIDR-GFP (green) together with a constitutive *glmS*::dsRed marker (magenta). The yellow line indicates the transect used to generate the fluorescence intensity profiles shown below the image; line profiles were quantified in Fiji (ImageJ). **B)** Phase-contrast and fluorescence images of *rne*(IDR_Cc_)-GFP cells expressing the chimeric *rne* (the *C. crescentus* IDR replacement, *rne*(IDR_Cc_)-GFP), as in (A). **C)** Colocalization between GFP and dsRed signals, expressed as Pearson correlation coefficients calculated from 30 cells from three biological replicates for *rne*(ΔIDR)-GFP and *rne*(IDR_Cc_)-GFP. Bars show mean values and error bars indicate standard deviation. (****, p < 0.0001; two-tailed t test with unequal variance). **D**) Demographs of GFP fluorescence intensity along the medial axis of individual *rne*-GFP cells (n = 171 cells), *rne*(ΔIDR)-GFP cells (n = 715 cells), and *rne*(IDR_Cc_)-GFP cells. Cells are ordered by increasing cell length; intensity is plotted along a normalized cell axis. **E)** Total GFP fluorescence intensity per cell (arbitrary units), reporting relative RNase E protein abundance, in *rne*-GFP cells, *rne*(ΔIDR)-GFP cells, and *rne*(IDR_Cc_)-GFP cells; bars show mean ± standard deviation (***, p < 0.001; two-tailed t test with unequal variance). **F)** Western blot of *rne*-GFP, *rne*(ΔIDR), and *rne*(IDR_Cc_) strains; molecular-weight markers (kDa) are indicated at left.

**Figure S3:**
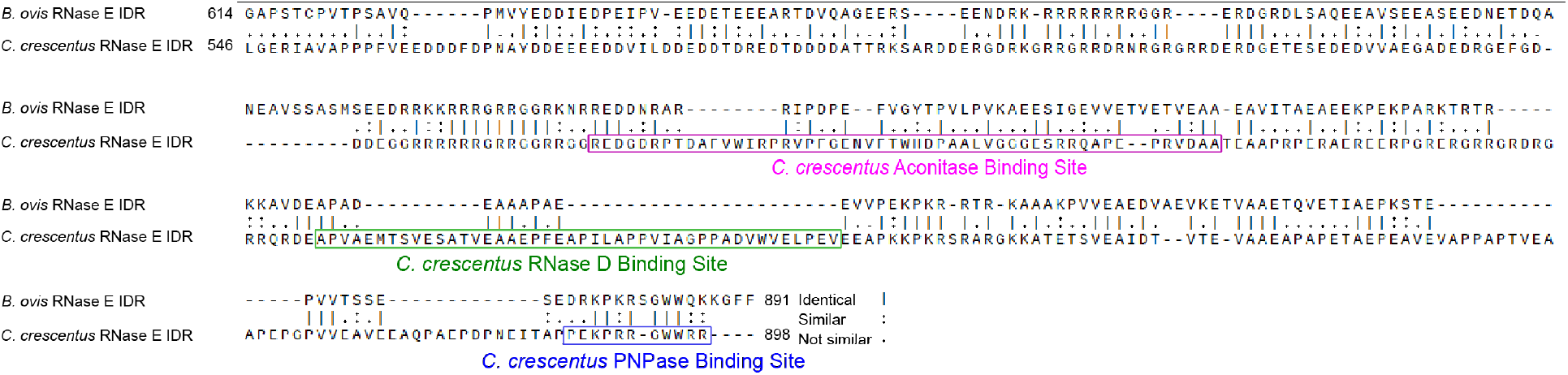
***B. ovis* RNase E IDR is highly divergent from the *C. crescentus* RNase E IDR.** The *B. ovis* RNase E IDR shares only 30.29% overall sequence identity with the *C. crescentus* RNase E IDR (Needleman-Wunsch global alignment, alignment matrix: BLOSUM62, gap open penalty:10.0, gap extend penalty: 1.0). Known *C. crescentus* degradosome binding sites within the IDR also show low sequence identity when aligned to the *B. ovis* IDR: RNase D binding site (40%), aconitase binding site (39%), and PNPase binding site (60%) (Smith-Waterman local alignment, alignment matrix: BLOSUM62, gap open penalty:10.0, gap extend penalty: 1.0).

**Figure S4:**
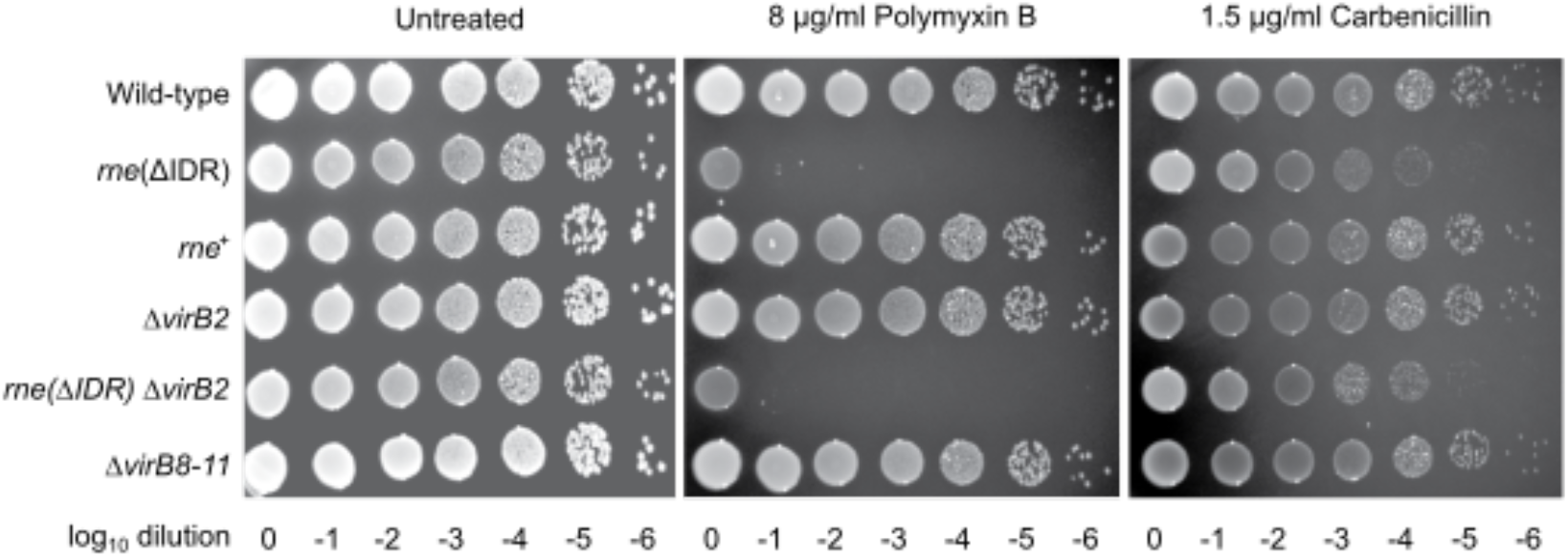
***virB* mutants do not impact stress sensitivity.** Spot dilution assays of wild type, *rne*(ΔIDR), *rne*⁺, Δ*virB2*, *rne*(ΔIDR) Δ*virB2,* and *ΔvirB8-11* for stress sensitivity. Strains on TSA blood agar were resuspended, normalized to OD_600_ = 0.3, serially diluted 10-fold, and spotted (5 µL) onto TSA blood agar plates lacking stressor or supplemented with polymyxin B (8 µg/mL), carbenicillin (1.5 µg/mL). Plates were incubated at 37°C and imaged after 72-96 hours. Dilution plating experiments were repeated at least three times (n=3) in technical triplicate, and one representative experiment is shown.

**Figure S5:**
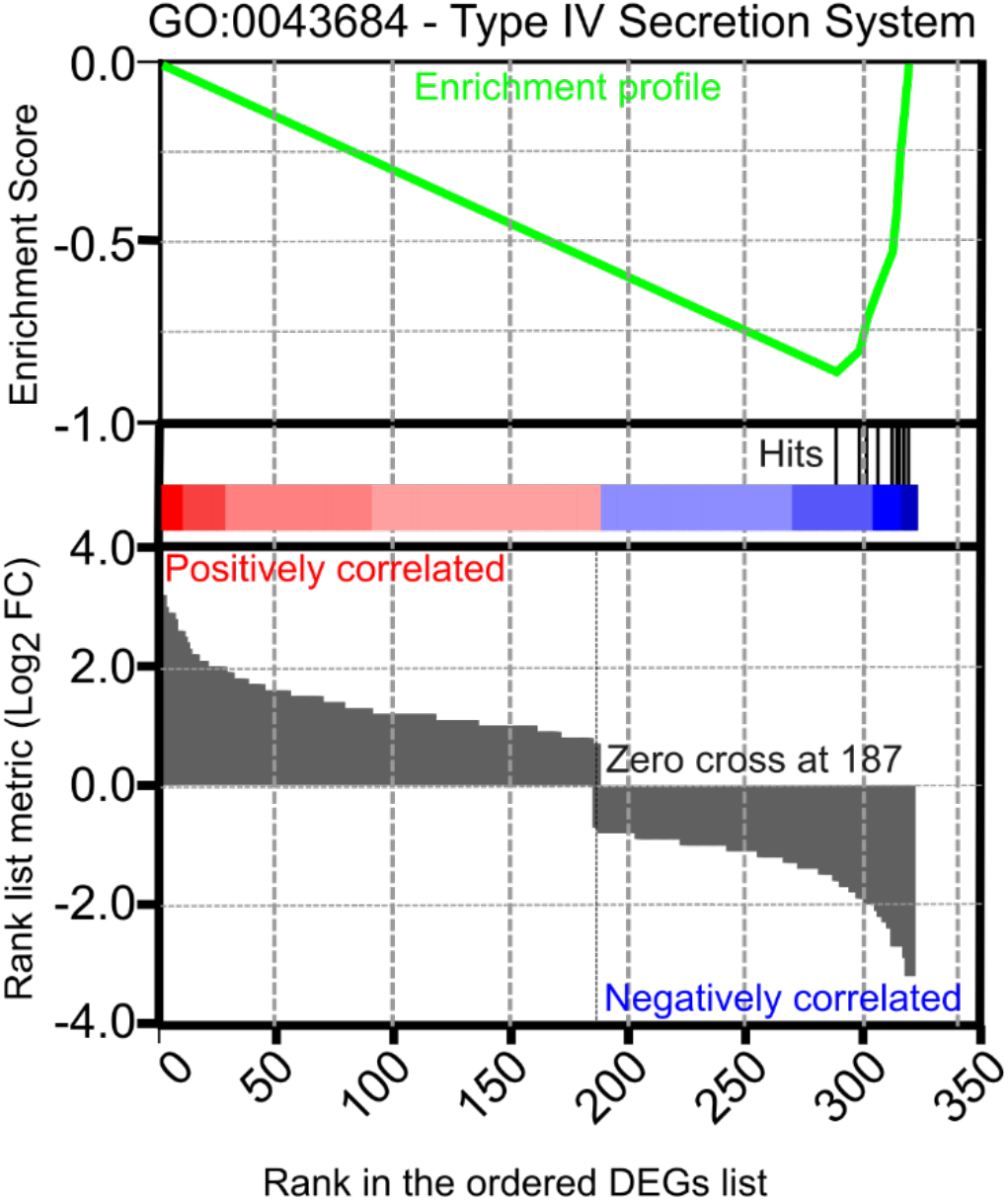
Annotated type IV secretion system (T4SS) components are enriched at the bottom of the ranked gene list and are collectively downregulated in the *rne*(ΔIDR) mutant. Gene Set Enrichment Analysis (GSEA) enrichment plot for GO:0043684 (type IV secretion system) across the ranked gene list. The green curve represents the running enrichment score, and vertical tick marks indicate the positions of genes annotated with the type IV secretion system GO term within the ranked list. The color bar and ranked-metric panel show the distribution of log_2_ fold changes used to rank the genes.

## Notes

### Competing Interest Statement

The authors have declared no competing interest.

